# Exploring MicroRNA Signatures in Pediatric Non-infectious Uveitis: Meta-Analysis and Molecular Profiling of Patient Samples

**DOI:** 10.1101/2024.09.27.615340

**Authors:** Olga Wawrzyniak, Dariusz Wawrzyniak, Michał Smuszkiewicz, Paweł Głodowicz, Anna Gotz-Więckowska, Katarzyna Rolle

## Abstract

**Purpose:** To find a distinct non-coding RNA characteristic for idiopathic uveitis in the pediatric population. To explore the autoimmune-related miRNA expression profile in pediatric patients with idiopathic uveitis (IU) and juvenile idiopathic arthritis-associated uveitis (JIA-AU) and find a common molecular background for idiopathic uveitis and other autoimmune diseases.

**Materials and methods:** The expression levels of miRNAs were analyzed by quantitative real-time PCR using serum samples from patients with idiopathic uveitis (n=8), juvenile idiopathic arthritis-associated uveitis (n=7), and healthy controls. We selected the most promising miRNAs from the original research papers: miR-16-5p, miR-26a-5p, miR-145-5p, miR-451a as markers for juvenile idiopathic arthritis, miR-23a-3p, miR-29a-3p, miR-140-5p, miR-193a-5p and miR-491-5p for uveitis in the adult population, and miR-125a-5p, miR-146a-5p, miR-155-5p, miR-223-5p, miR-223-3p characteristic for both diseases and confirm their expression changes in serum from children with idiopathic uveitis. We comprehensively reviewed the literature enrolling the papers that met the inclusion criteria (miRNA and non-infectious uveitis/juvenile idiopathic arthritis) and performed target prediction analysis of appoint miRNAs. It additionally confirmed that altered miRNAs target the immunologically involved genes.

**Results:** Immunological involved miRNAs such as miR-146a-5p and miR-155-5p show diverse expression levels in different patients as they interact with multiple targets. miR-204-5p is downregulated in both patient groups compared to healthy controls.

**Conclusion:** miR-204-5p and miR-155-5p are candidates for molecular markers of autoimmune uveitis. We did not identify the miRNAs specific only to idiopathic uveitis, but for the first time in the pediatric population, we confirmed that this disease entity shares a molecular basis with other autoimmune diseases. Further studies are required to elucidate the molecular interactions among miRNAs, cytokines, and transcription factors within the intricate immune response, particularly in the eye.

## 1. Introduction

Uveitis is the inflammation of the uveal membrane of the eye. It can be caused by infectious pathogens or be autoimmune or idiopathic. Pediatric patients represent 5-16% of all patients with uveitis [1,2]. The incidence and prevalence differ between countries and populations and there are about 4-5 new cases per 100 000 children in developed countries [3,4]. Although it can be categorized as a rare disease, it remains an important cause of irreversible blindness [5,6]. According to the Standardization of Uveitis Nomenclature (SUN) criteria, uveitis description includes onset, duration, course of inflammation, and anatomical localization [7]. 95% of uveitis in children is noninfectious with idiopathic uveitis (IU) and juvenile idiopathic arthritis-associated uveitis (JIA-AU) as the most common [2,8]. It is usually insidious, chronic, and persistent in duration [9].

Due to often delayed diagnosis, difficult examination, and limited treatment options in young patients, pediatric uveitis can lead to severe chronic inflammation, a higher risk of developing ocular complications, and permanent vision loss [10,11]. Major undesirable outcomes of the disease and its treatment are posterior synechiae, secondary cataract, secondary glaucoma, hypotony, and maculopathy. All can cause amblyopia [12]. Furthermore, the long-term effects of this condition and the use of topical and systemic medications over a long period impact vision-related quality of life and function [13]. Despite the improving guidelines for the management of patients with autoimmune diseases, pediatric uveitis remains a vision-threatening condition [12].

Juvenile idiopathic arthritis is the most frequent rheumatic disease in childhood, with an annual incidence between 8 and 22,6 per 100 000 children and prevalence rates between 70 and 401 per 100 000 children [14]. The diagnosis merges the group of conditions with a common feature: arthritis. The International League of Associations for Rheumatology (ILAR) criteria for the classification of JIA distinguish 7 subtypes of this disease. Oligoarticular arthritis and polyarticular arthritis constitute about 45% of all cases and are strongly associated with asymptomatic uveitis in nearly 45% of patients, especially with the presence of antinuclear antibodies (ANA) [15]. Acute symptomatic uveitis usually correlates with enthesitis-related arthritis and HLA-B27 occurrence [16]. Different subtypes of JIA are often considered to be the childhood onset of rheumatological diseases in adults. Polyarticular RF+ JIA usually develops into rheumatoid arthritis (RA), and enthesitis-related arthritis with positive HLA-B27 status passes into ankylosing spondylitis (AS) [16].

Pediatric patients diagnosed with juvenile idiopathic arthritis are under ophthalmological supervision, so the early signs of uveitis can be spotted and properly treated. Also, the general treatment of rheumatoid disease helps with maintaining ocular inflammation. Although several guidelines define the current and most efficient course of follow-up in JIA-AU patients, a group of children with idiopathic uveitis or with uveal inflammation before the rheumatoid diagnosis remains challenging for clinicians [17,18].

Noncoding RNAs are a heterogeneous group of small RNA particles: microRNAs (miRNAs), long-non-coding RNAs, and circular RNAs. All regulate gene expression at the posttranscriptional level controlling different cellular mechanisms [19]. Many miRNAs identified in humans are conserved in other animals. These short, ∼22 nt long particles interact with mRNA and influence developmental processes and diseases [20]. Loss-of-function studies disrupting miRNA genes in mice and rats have revealed their involvement in immunological processes, such as the increased proliferation of progenitors and hyperactive neutrophils and macrophages, inflammatory cytokine production, or Th17 cell activation [21,22]. Changes of the miR-93, miR-155, miR-146a, miR-182, and miR-223 expression levels modify numerous protein transcription for example Signal transducer and activator of transcription 3 (STAT3), interleukin-1 receptor-associated kinase 4 (IRAK4), or Forkhead box O3 (FOXO3) involved in immune reaction in induced non-infectious uveitis [23–28]. In cardiovascular and neurological pathologies and oncological processes, the participation of miRNA is well explored [29–31]. In autoimmune diseases, they are increasingly attracting attention as potential markers of disease activity and as targets for therapy [32]. In addition to miRNA, lncRNA, and circular RNA also play a role in immunological pathways by interacting with mRNA, acting as decoy particles, or producing proteins with similar or different activity to their main protein [33–35].

Detailed information about non-coding RNA genesis and function in ocular disorders authors presented in previous publications [35,36].

In the present study, we determined an expression profile pattern of inflammatory microRNAs in the blood of pediatric non-infectious uveitis patients and compared it with previous studies performed on samples from adult patients. We also summarized for the first time the molecular background and revealed the similarities and differences in miRNAs expression levels between subgroups of non-infectious uveitis in children. We highlighted a set of microRNAs that may have a regulatory function in pediatric uveitis. Thus, we have identified a mechanism caused by altered expression of miRNAs that may be related to these diseases, providing new insights into potential diagnostic and therapeutic targets.

## 2. Materials and methods

### 2.1 Patients

This study was approved by the Bioethics Committee at the Poznań University of Medical Sciences (NR 951/16), and informed consent was obtained from all parents/legal guardians of the patients, and patients above 16 years old.

The study included 15 patients with non-infectious uveitis and 3 healthy controls between 6 and 18 years old, admitted to the Pediatric Ophthalmology Clinic of Poznań University of Medical Sciences between 2019-2022.

The epidemiological data of the patients are summarized in **Table 1**. Additional information on the individual patients is summarized in **Supplementary Table S1**.

**Table 1.** Clinical characteristics of the cohorts investigated in this study.

|  | **JIA-AU** | **IU** | **HC** |
| --- | --- | --- | --- |
| **Number of patients** | 7 | 8 | 3 |
| **Age at the diagnosis, mean ± SD** | 6.4 ± 4 | 12.8 ± 2.9 | N/A |
| **Age at sample collection, mean ± SD** | 14 ± 3.2 | 15 ± 2 | 8.7 ± 1.5 |
| **Male/female** | 3/4 | 5/3 | 1/2 |
| **ANA +, (number of tested patients)** | 7 (7) | 1 (5) | N/A |
| **HLA-B27 +, (number of tested patients)** | 1 (2) | 0 (5) | N/A |

JIA-AU – juvenile idiopathic arthritis-associated uveitis, IU – idiopathic uveitis, HC-healthy control, N/A – not applicable.

### 2.2 Extraction of total RNA and DNase treatment

The patient’s blood samples were diluted at a 1:1 ratio in phosphate-buffered saline (PBS). Subsequently, 750 µl of TRI Reagent^®^ Solution was added to 250 µl of the blood dilution, followed by a 5-minute incubation period at room temperature. The lysate was gently mixed to ensure thorough homogenization. 200 µl of chloroform was added to the TRIzol-blood mixture, followed by an incubation period of 10 minutes with repeated mixing to facilitate phase separation. The aqueous and organic particles were separated by centrifugation for 15 minutes at 13000 rpm at 4°C. The supernatant containing the RNA was carefully transferred to a new Eppendorf tube, and isopropanol was added to the supernatant at an equivalent volume to precipitate the RNA. The samples were incubated overnight at –20°C or for 3 hours at –80°C and centrifuged at 13000 rpm for 30 minutes at 4°C to pellet the RNA. The RNA pellet was washed with 1 ml of cold 80% ethanol and centrifuged at 13000 rpm 4°C for 10 minutes. After removal of the ethanol, the RNA pellet was air-dried for 10 minutes at room temperature and resuspended in 20 µl of double-distilled, sterile water. The quality of extracted RNA was measured by NanoDrop 2000 spectrophotometer and confirmed by the agarose gel electrophoresis. Afterward, RNA samples were subjected to DNase I treatment using a ready-to-use DNA-free™ Kit DNAse Treatment & Removal (Thermo Fisher Scientific).

### 2.3 Polyadenylation reaction, miRNA reverse transcription, and quantitative RT-qPCR

The reverse transcription for miRNA was performed in two steps. First, 500 ng of total RNA was polyadenylated by using *E.coli* Poly (A) polymerase, followed by reverse transcription for miRNA using AffinityScript One-Step RT-PCR Kit (Agilent Technologies) with an anchored-oligo(dT) primer, both according to the manufacturer’s protocol. Complementary DNA (cDNA) was used as a template in real-time quantitative reverse transcription PCR (qRT-PCR), with the use of CFX96 Real-Time System thermal cycler (BioRad), in three technical replicates. Relative expression was analyzed in the CFX96 Real-Time System thermal cycler (BioRad) Software version 2.0. In the case of miRNA normalization, the mean level of ACTN1, GAPDH, HPRT, SDHA as an endogenous control was used. A list of sequences of primers 5’-3’ are presented in the **Supplementary Table S2**. Statistical significance was calculated using unpaired t test with Welch’s, correction, * *p* ≤ 0.05, ** *p* ≤ 0.01, *** *p* ≤ 0.001, **** *p* ≤ 0.0001. Bars representing not significant results (*p* > 0.05), are not shown on the graphs.

### 2.4 Selection of candidate microRNAs

The 19 microRNAs tested by qRT-PCR in blood samples were selected from a literature review. This literature research was performed in 2019 on the PubMed database and focused on determining a list of microRNAs with potential or known functions in idiopathic uveitis disease, as well as in autoimmune diseases. This biased approach to select candidate microRNAs was intentionally used to highlight potential microRNAs involved in inflammation, the main biological feature of uveitis.

### 2.5 Meta-analysis

To better organize our work we follow the Preferred Reporting Items for Systematic reviews and Meta-Analyses (PRISMA) 2021 guidelines [37]. Literature searches were conducted using the PubMed database. Considering there was only one publication on noncoding RNAs and pediatric uveitis, we expanded the search strategy. The combinations of search terms included ‘micro/mi RNA’, ‘non-coding RNA’, ‘pediatric uveitis’, ‘uveitis in children’, ‘uveitis’ “autoimmune uveitis’ ‘noninfectious uveitis’, and as pediatric uveitis mostly coexists with juvenile idiopathic arthritis also with ‘juvenile idiopathic arthritis’.

Original research papers on miRNAs in non-infectious uveitis and juvenile idiopathic arthritis that met the inclusion criteria were thoroughly examined. The inclusion and exclusion criteria are summarized in **Table 2**.

**Table 2.** Inclusion and exclusion criteria.

| Inclusion criteria | Exclusion criteria |
| --- | --- |
| original paper | review |
| human samples | animal models |
| idiopathic uveitis, autoimmune uveitis,<br>juvenile idiopathic arthritis | infectious uveitis |
| miRNA expression level | SNPs |

### 2.6 Prediction of miRNA’s targets

Target annotation analysis and network visualization were performed using the R environment (version 4.4, https://www.r-project.org) and appropriate packages according to corresponding reference manuals. Identification of miRNA/gene regulatory interactions was performed *in silico* between selected miRNAs and uveitis and juvenile arthritis-associated genes harvested from DisGeNET 7.0 database (https://www.disgenet.org/) [38–40], using multiMiR 1.26.0 package (https://bioconductor.org/packages/release/bioc/html/multiMiR.html) [41]. The analysis included the identification of both experimentally validated (miRecords, miRTarBase, and TarBase databases) and predicted (DIANA-microT-CDS, ElMMo, MicroCosm, miRanda, miRDB, PicTar, PITA, and TargetScan databases) miRNA/gene interactions. The regulatory network visualization with obtained interactions was performed using Cytoscape v3.10.2 software (https://cytoscape.org/) and MiRTargetLink 2.0 [42,43].

### 2.7 Enrichment analysis

Pathway enrichment analysis was performed using the ToppGene Suite (https://toppgene.cchmc.org/enrichment.jsp), where a list of selected miRNAs and a list of genes with which they interact were introduced [44]. The default *Homo sapiens* genome was used as a background. Terms of the Kyoto Encyclopedia of Genes and Genomes (KEGG) pathways [45], REACTOME [46], and Gene Ontology (GO) categories [47] were searched.

### 2.8 Statistical analysis

Results at *p* < 0.05 were considered statistically significant. Statistical analyses were performed in GraphPad Prism 8.0.1 (GraphPad Software, San Diego, CA, USA). The analyses included the Shapiro-Wilk test (to assess the compliance of the examined variables with the normal distribution), the student’s *t*-test for variables on a quantitative scale (data presented as an average). GraphPad Outlier calculator (https://www.graphpad.com/quickcalcs/grubbs1/) was used to determine the significance of the most extreme values. Significant outliers, *p* < 0.05 were not shown on the graphs (**Supplementary Table 3**).

## 3. Results

### 3.1 miRNA expression profile in blood samples

To establish the molecular profile of miRNAs, we analyzed the expression of 19 preselected microRNAs in the 18 blood samples from uveitis (15) and healthy (3) patients by qRT-PCR (**Figure 1-2**). The expression level of selected miRNAs in juvenile idiopathic arthritis-associated uveitis (JIA-AU) patients is not identical in all subjects (**Figure 1**). This may be because this group includes patients with different subtypes of JIA who are treated with immunosuppressive and immunomodulatory drugs in various combinations. For patients with JIA-AU (5 upregulated and 14 downregulated) and for patients with IU (13 upregulated and 6 downregulated) miRNAs were examined compared with healthy controls. Among these miRNAs, 4 upregulated miRNAs (miR-182-5p, miR-223-3p, miR-23a-3p and miR-30b-5p) and 5 downregulated miRNAs (miR-125a-5p, miR-135a-5p, miR-135b-5p, miR-193a-5p and miR-204-5p) were common to the JIA and IU. The rest of studied miRNAs, such as miR-140-5p, miR-145-5p, miR-146-5p, miR-155-5p, miR-16-5p, miR-26a-5p, miR-29a-3p, miR-451a, miR-491-5p, miR-223-5p, show an opposite pattern in JIA-AU and IU (**Figure 2**).

**Figure 1.**
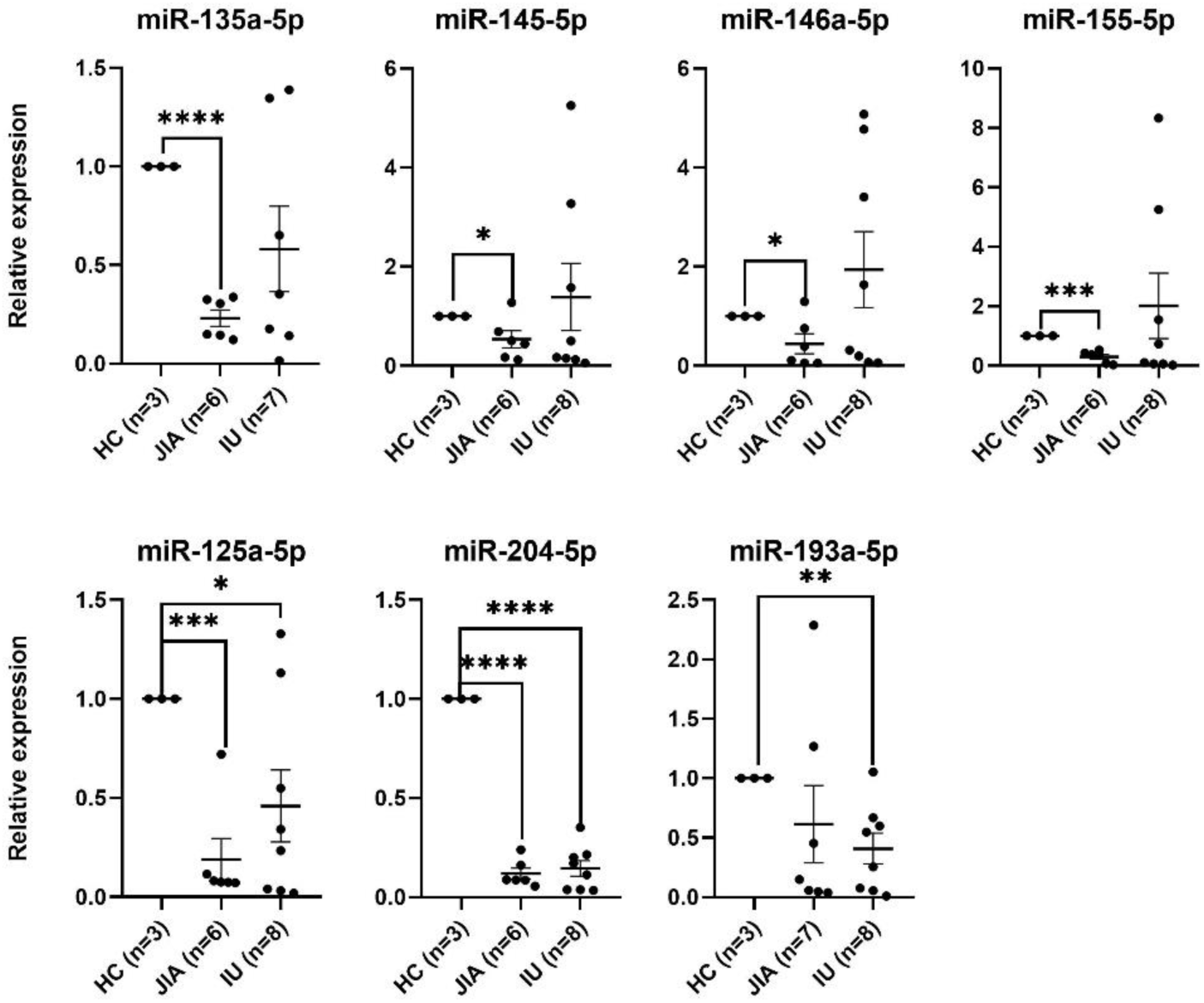
Differentially expressed miRNAs using qRT-PCR between JIA-AU and IU patients and healthy donors. n represents number of patients without significant outliers. Statistics was performed with unpaired t test with Welch’s, correction, * *p* ≤ 0.05, ** *p* ≤ 0.01, *** *p* ≤ 0.001, **** *p* ≤ 0.0001. Bars representing not significant results (*p* > 0.05), are not shown on the graphs.

**Figure 2.**
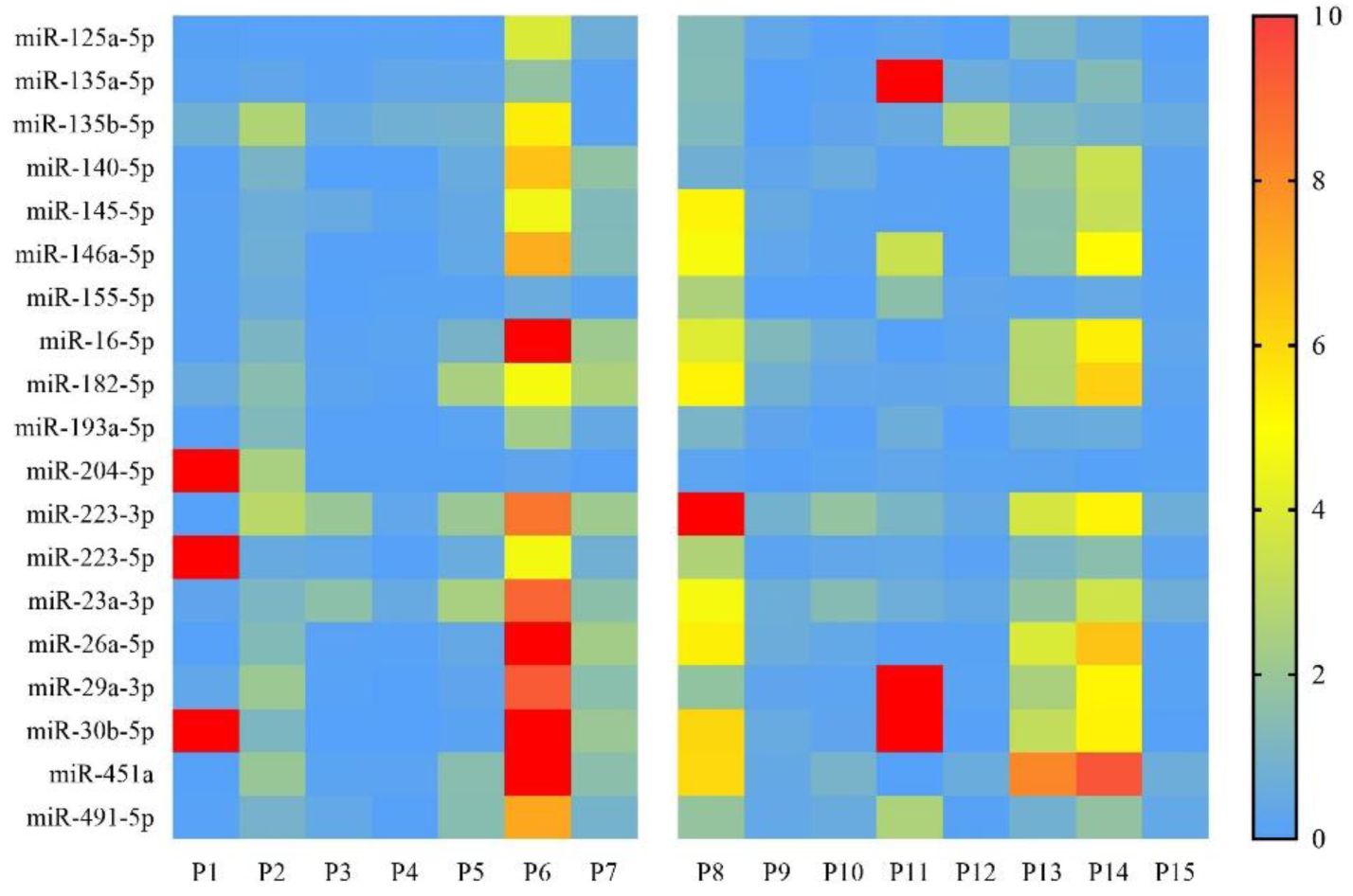
Heatmap showing differential miRNA expression in diseased *vs* normal control group. The left panel represents JIA-AU patients and right panel represents IU patients. The green colour indicates lower than mean intensity, and red represents higher than mean intensity. Each row represents a miRNA and each column represents a patient’s sample (*p* < 0.05, logFC > |1|).

### 3.2 Meta-analysis

One of the aims of the present study was to perform a meta-analysis of miRNA expression profiling studies in human uveitis samples to complement and compare data from RT-qPCR analysis performed on samples from children suffering from uveitis. One hundred fifty-two papers were obtained using combinations of the entries described above. Of these, 68 met the inclusion criteria listed in **Table 2**. 35 original research papers were included in the comprehensive review, 17 on juvenile idiopathic arthritis and 18 on non-infectious uveitis. Our search included English-language publications from 2000 to November 2023. **Figure 3** based on PRISMA 2021 guidelines shows the literature selection process.

**Figure 3.**
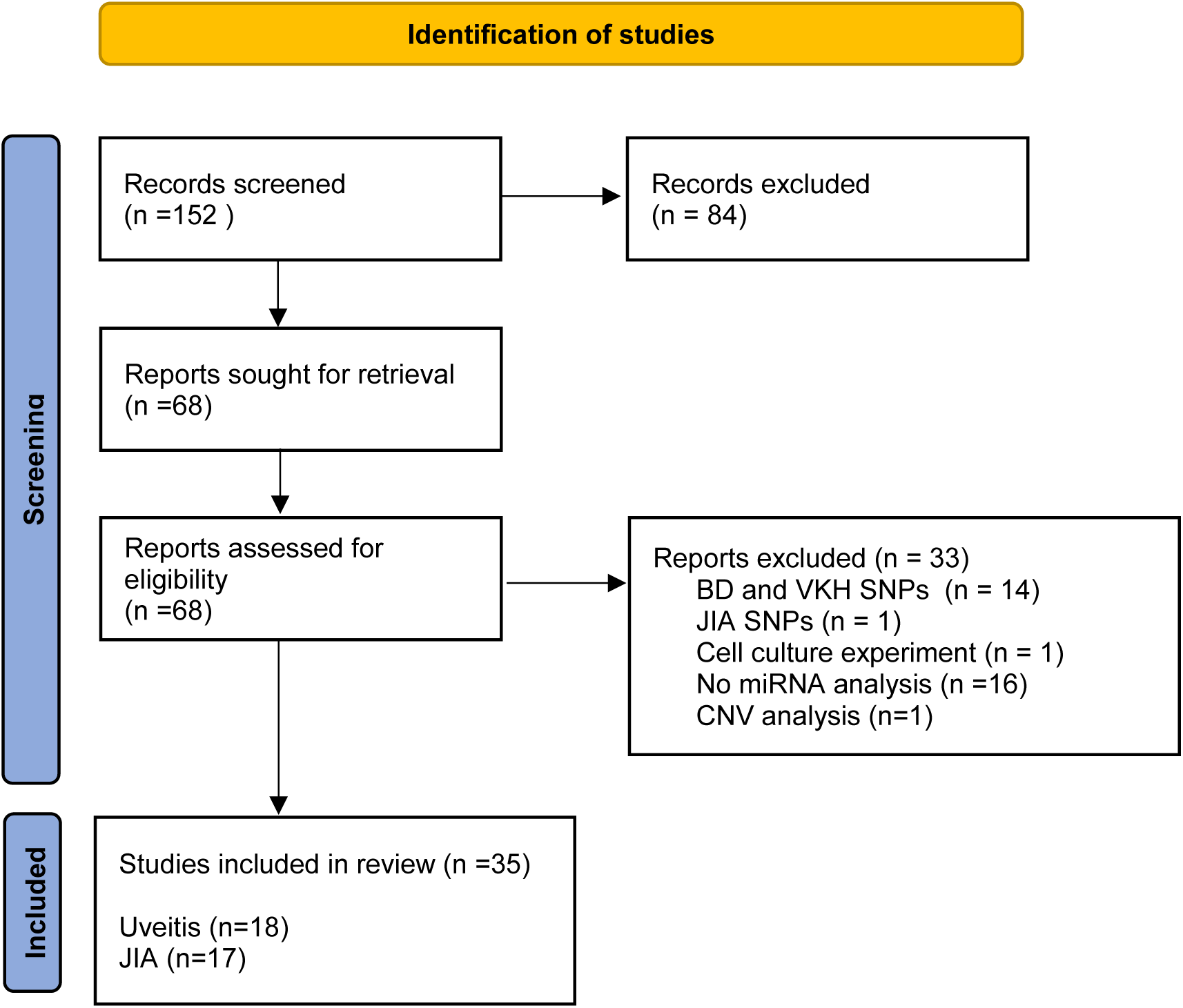
Identification of the studies included in the meta-analysis (search process based on PRISMA 2021).

#### 3.2.1 miRNA in non-infectious uveitis

The potential use of microRNAs as markers and predictors of specific types of uveitis has been best explored in Behcet disease (BD) and Vogt-Koyanagi-Harada disease (VKH) [48–52]. These two subtypes of uveitis are rare in children. Behcet disease in pediatric patients mostly starts in late childhood and a full diagnosis with all criteria met is usually possible after age 16 [53,54]. It is common in Turkey, Japan, China, Saudi Arabia, Israel, and Iraq but rare in most developed countries except for the Mediterranean [55]. The well-explored fields in BD and VKH are miR-146a SNPs as well as miR-196a2 and miR-182 [48,52,56–61]. As we focused on pediatric patients and miRNA expression levels, we excluded research papers on SNP variants in BD and VKH.

The only study that includes pediatric patients with uveitis and miRNAs explores the miR-146a SNPs in the Han Chinese population. In the group of 520 patients and 1204 healthy controls included in the study, the researchers found that the frequencies of the rs2910164 CC genotype and the C allele of miR-146a were significantly decreased in patients compared to healthy controls. This suggests that miR-146a may be a susceptibility factor for pediatric uveitis. They also correlated the SNP rs10893872 of Ets-1, an oncogene and transcription factor, with uveitis predisposition. The CC genotype and C allele frequencies of this SNP were significantly increased in patients compared to controls. Ets1 can bind to the miR-146a promoter region and significantly affect miR-146a activity [62].

To extract the most promising noncoding RNAs involved in noninfectious ocular inflammation we had to extend our research to adult patients in this field. Among 19 studies performed on human samples, only 5 were conducted on serum, 1 on vitreous, and the rest on peripheral blood mononuclear cells (PBMCs). Most papers explore miRNA expression in Behcet disease or combination with Vogh-Harada-Koyanagi disease. Only 4 studies investigate other types of uveitis: HLA-B27-associated acute anterior uveitis (AAU), idiopathic intermediate uveitis (IIU), anterior uveitis associated with ankylosing spondylitis or non-infectious posterior and panuveitis [63–66]. Verhagen and colleagues discovered and independently confirmed a group of microRNAs associated with noninfectious uveitis (NIU). miR-140-5p, miR-491-5p, miR-223-3p, miR-223-5p, miR-193a-5p, miR-29a-3p, and U6 snRNA showed strong correlation, prompting researchers to treat them as single miRNA cluster associated with uveitis. Six of these miRNAs were found to target a total of 37 genes. Analysis of these common gene targets revealed significant involvement in pathways related to inflammation and ocular biology [66].

Research on miR-146a expression in peripheral blood mononuclear cells (PBMCs) from Behçet’s disease (BD) and acute anterior uveitis (AAU) patients has yielded mixed results. Three studies in BD patients and one in AAU patients found reduced miR-146a expression, consistent with its potential anti-inflammatory role [51,67–69]. However, O’Rourke and colleagues observed increased levels of miR-146a, suggesting that it acts as a regulator of the immune response, in contrast to the pro-inflammatory activity of miR-155. These findings suggest a possible feedback mechanism between the pro-inflammatory miR-155 and the regulatory miR-146a, mediated by Toll-like receptor (TLR) signaling [65]. Similarly, miR-155 expression is usually upregulated in most studies conducted in PBMCs from BD and AAU patients [50,65,68]. Yet, Zhou et al. showed that miR-155 is downregulated in PBMCs and CD4+ cells in the active phase of BD [57]. In addition, Ahmadi et al. reported that both miR-155 and miR-146a expression were decreased in BD patients, which may play a role in the reduced number of Treg cells noted in BD patients [67]. This evidence supports the idea that miRNA expression may fluctuate during different stages of the disease and possibly in response to treatment.

MiRNAs have also been investigated for their diagnostic potential in intraocular lymphoma and uveitis [70]. Expression profiles of miR-155, miR-200, and miR-22 were found to distinguish between the two diseases. **Table 3** summarizes papers about noninfectious uveitis enrolled in this study.

**Table 3.** miRNA in non-infectious uveitis. BD: Behcet Disease; VKH: Vogt-Kyanagi-Harada disease, HC: healthy control, AAU: acute autoimmune uveitis, IIU: idiopathic intermediate uveitis, BU: Birdshot Uveitis, SS: suspected sarcoidosis, PPU: panuveitis/posterior uveitis, S: Sarcoidosis, AIU: anterior idiopathic uveitis, AAU AS+/-: Acute anterior uveitis with or without ankylosing spondylitis, PBL: primary B-cell lymphoma, PBMC: peripheral blood mononuclear cells.

| Patients and diagnosis | miRNA up-regulated | miRNA down-regulated | Sample | miRNA identification method | Reference |
| --- | --- | --- | --- | --- | --- |
| patients n = 54,<br>HLA-B27 AAU (19)<br>IIU (15)<br>BU (19)<br>HC n = 26 | miR-140-5p, miR-491-5p, miR-223-3p, miR-223-5p, miR-193a-5p, miR-29a-3p U6 snRNA- uveitis associated cluster | - | serum | Open Array (TaqMan), RT-qPCR (TaqMan) | Verhagen FH. et al., 2018 [66] |
| patients n = 40<br>BD (10),<br>Sarcoidosis (17);<br>VKH (13)<br>HC n = 11 | miR-1226-3p, miR-133b, miR-193b-3p, miR-204-5p, miR-2277-5p, miR-326, miR-422a, miR-4652-3p, miR-4698, miR-483-3p, miR-495-5p, miR-519d-5p, miR-520f-5p, miR-542-5p, miR-548ab, miR-6734-3p, miR-6744-3p, miR-6769b-3p, | miR-1301-5p, miR-193a-5p, miR-300, miR-302c-5p, miR-3605-5p, miR-3612, -miR-379-5p, miR-3936, miR-3978, -miR-4461, miR-4633-3p, miR-4681, miR-4694-3p, miR-4724-5p, miR-4755-3p, -miR-5190, miR-5192, miR-6501-3p, hsa-miR-6728-5p, miR-6804-5p, miR-6817-5p, miR-6823-5p, miR-6853-5p, miR-7162-3p, miR-758-5p, miR-8060, miR-920, | serum | Microarray (3D-Gene miRNA Labeling kit; 3D-Gene Human miRNA Oligo Chip) | Asakage M, et al. 2020 [63] |
| patients n = 15,<br>HLA-B27 + (7)<br>AIU ( 8)<br>HC n = 5 | miR-146a, miR-155, miR-125a-5p | - | PBMC | TaqMan miRNA Assays | O'Rourke M, et al. 2019 [65] |
| patients n = 21<br>PPU/S (11)<br>SS (10)<br>HC n = 16 | miR-455-3p, let-7b-5p, miR-5000-5p, miR-6716-3p, miR-6806-3p, miR-106a-5p, miR-4663, miR-548a-3p, miR-450b-5p, miR-17-5p, let-7c-5p, miR-5189-3p, miR-6716-3p, | miR-6805-5p, miR-22-3p, miR-23a-3p, miR-1184, miR-425-5p, miR-3621, miR-4467, miR-191-5p, miR-3613-3p, miR-4668-5p, miR-4674, miR-185-3p, miR-378h, miR-3197, | serum | Microarray (GeneChip® miRNA 4.0 arrays) | Saito S, et al. 2022 [64] |
| patients n = 29<br>PBL (17)<br>IU (12) | miR-155, miR-200, miR-22 | - | vitreous | Quantitive real-time PCR-based miRNA panel (StepOnePlus RT-PCR System) | Tuo J, et al. 2014 [70] |
| patients n = 62<br>BD (active – 23, inactive – 19)<br>VKH (21)<br>HC n = 20 | - | miR-155 in active BD | PBMC<br>Dendritic cells<br>CD4+ cells | miRNA TaqMan assay | Zhou Q, et al. 2012 [57] |
| patients n = 56<br>BD (28)<br>Neuro BD (28)<br>HC n = 30 | - | miR-185 in BD and neuro BD | PBMC | RT-qPCR (miScript II RT Kit, miScript Primer Assay) | Ugurel E, et al. 2015 [79] |
| patients n = 6<br>BD | miR-4505, miR-149-3p | let-7d-5p, miR-181a-5p, miR-146a-5p, miR-361-5p, miR-532-3p, miR-423-5p, miR-200c-3p, miR-30e-5p, miR-28-5p, miR-30c-5p, miR-330-3p, miR-194-5p, miR-423-3p, miR-28-3p, miR-15b-5p, miR-30d-5p, miR-193a-5p, miR-192-5p, miR-152-3p, miR-25-3p, miR-181d-5p, let-7f-5p, miR-92b-3p, miR-30a-5p, miR-223-3p, miR-505-3p, miR-128-3p, miR-148b-3p, miR-328-3p, miR-195-5p, let-7e-5p, miR-29b-1-5p, miR-628-3p, miR-92a-1-5p, miR-27b-3p, miR-671-3p, miR-151a-3p, miR-486-5p, miR-199a-3p, miR-199b-3p, miR-126-3p, miR-584-5p, miR-199a-5p, miR-139-5p, miR-143-3p | PBMC | Microarray (Affymetrix GeneChip® miRNA 4.0)<br>Real time PCR (TaqMan® Advanced miRNA assay) | Puccetti, A. et al.<br>2018<br>[51] |
| patients n = 46<br>BD<br>HC n = 26 | miR-155, |  | PBMC | Real-time PCR (ABI Prism 7000 Sequence Detection System) | Na SY, et al.<br>2016<br>[50] |
| patients n = 47<br>BD<br>HC n = 61 | miR-155 | miR-146a | PBMC | RT- PCR | Kolahi S, et al.<br>2018<br>[68] |
| patients n = 46<br>BD<br>HC n = 70 | - | miR-21 | PBMC | qRT-PCR (ExiLent SYBR Green Master mix) | Jadideslam G, et al.<br>2019<br>[80] |
| patients n = 47<br>BD<br>HC n = 34 | miR-571, miR-127, miR-573, miR-770, miR-370, miR-139-3p, miR-720, miR-330 | miR-200b, miR-520c-3p, miR-199a-3p, miR-224, miR-98 | PBMC | Profiling miRnome (Human MicroRNA TaqMan Low-Density Array) | Erre GL, et al.<br>2015<br>[81] |
| patients n = 46<br>BD<br>HC n = 48 | miR- 224- 5p, miR- 206, miR- 653- 5p | - | serum | Microarray (Agilent Human miRNA 8×15 k Microarray kit v3.0 and miRNA Complete Labelling and Hyb Kit)<br>PTA-PCR (miRNA- specific primers) | Emmi G, et al.<br>2022<br>[82] |
| patients n = 51<br>BD<br>HC n = 45 | miR - 181b | - | serum | RT-PCR (miScript miRNA PCR custom array) | El Boghdady NA, et al.<br>2019<br>[83] |
| patients n = 38<br>VKH<br>HC n = 25 | - | miR- 20a-5p | PBMC | Quantitative real-time PCR (miDETECT A Track Kit, ABI 7500 System) | Chang R, et al.<br>2018<br>[84] |
| patients n = 47<br>BD<br>HC n = 58 | miR-25, miR-106b, miR-93, miR-326 | miR - 155, miR-146a | PBMC | RT-PCR (LightCycler 2.0 RT-PCR System) | Ahmadi M, et al. 2019 [67] |
| patients n = 786<br>AAU AS+ (384);<br>AAU AS- (384)<br>HC n = 660 | miR 9-3 in high CNV group | miR-9-3, miR-143, miR-146a | PBMC | TaqMan-based qPCR | Yang L, et al. 2017 [69] |
| Patients n = 19<br>Initial onset VKH = 6<br>Recurrent VKH = 6<br>HC = 6 | miR-206 in both stages of the disease | miR-7704 in recurrent stage | PBMC | NEBNext® Multiplex Small RNA Library Prep Set for Illumina; TruSeq SR Cluster Kit v3-cBot-HS; Illumina Hiseq 2500/2000 platform | Gou K. et al. 2022 [85] |

#### 3.2.2 miRNA in juvenile idiopathic arthritis

There are multiple differences between the selected studies regarding clinical characteristics, type of tissue examined, and miRNA isolation and analysis method. However, changes in the expression of miR-16, miR-125, miR-146a, miR-155, and miR-223 are confirmed. In juvenile arthritis, miR-146a is up-regulated in PBMCs, synovial fluid, and serum. The increased expression of miR-146a occurs in response to inflammatory processes. This microRNA promotes the alternative activation of M2 macrophages, associated with regulatory and anti-inflammatory functions. Additionally, miR-146a inhibits the polarization of macrophages towards the M1 phenotype [71–73].

The expression of miR-155 shows inconsistency across different studies. It regulates the immune response pathway by modulating IL2 and IL6 expression and can be involved in apoptosis and granulopoiesis. Research by Lashine and colleagues suggests its potential as a biomarker to distinguish lupus-associated arthritis from other inflammatory arthritic conditions. Their results showed reduced miR-155 expression in PBMCs from patients with lupus-associated arthritis, and increased expression in individuals with juvenile arthritis [74]. MiR-155 was also reported to be elevated in synovial fluid (along with miR-146a and miR-6764-5p) from patients with JIA compared to septic arthritis (SA). Overexpression of these three miRNAs in SF distinguishes JIA from septic arthritis and suggests their role in autoimmunity rather than infectious inflammation [72].

Ma et al. found decreased serum levels of miR-155 in poly-JIA patients compared to controls. Conversely, Demir et al. observed significantly elevated miR-155 levels in both active and inactive JIA. Despite these conflicting findings, both studies agree on the increased expression of miR-16 in JIA patients compared to healthy controls. Furthermore, plasma miR-16 levels have been shown to increase during active JIA and remain elevated in remission [71,75].

In contrast to miR-16, miR-223 appears to correlate with disease activity. Elevated levels of miR-223 have been observed during active phases of both systemic and polyarticular JIA, suggesting its potential as a biomarker of disease activity [76]. Levels of miR-300, miR-144, miR-133b, and miR-744, which are correlated with leukocyte adherence and transmigration pathways, differ between treated and untreated patient groups. It can be a useful tool in assessing the effectiveness of the treatment [77]. It is also reported, that miRNAs expression profile can change locally to promote joint-restricted inflammatory reactions. miR-23a regulates mitochondrial reactive oxygen species (ROS) release and is overexpressed in synovial fluid mononuclear cells compared to peripheral blood mononuclear cells. [78]. **Table 4** summarizes papers on the subject of miRNA in juvenile idiopathic arthritis enrolled in this study. **Figure 4** illustrates common and differentiating miRNAs for idiopathic uveitis and other autoimmune conditions.

**Figure 4.**
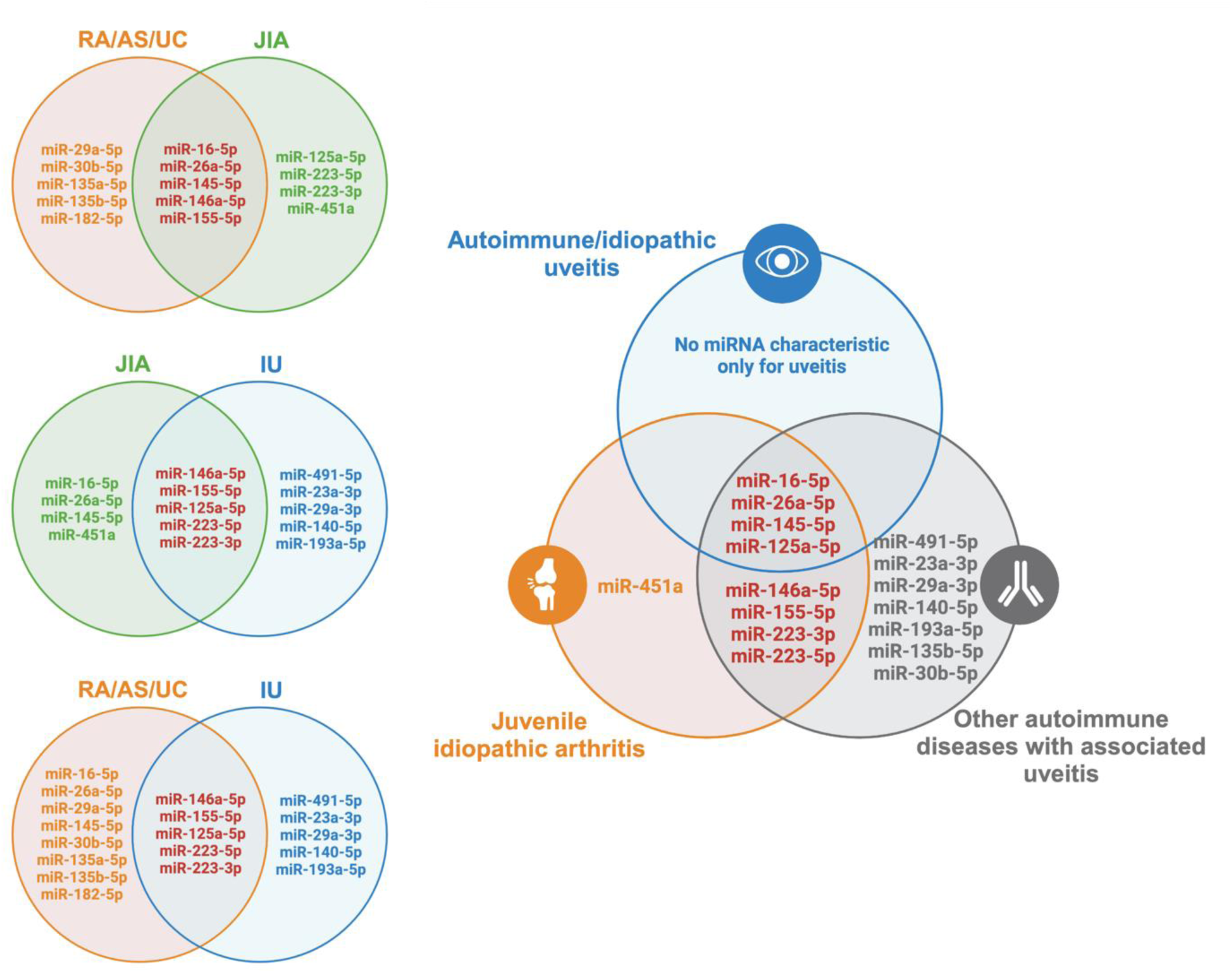
Selected miRNAs involved in uveitis, juvenile idiopathic arthritis, and other autoimmune diseases. Changes in the expression of miR-125a-5p miR-146a-5p miR-155-5p miR-223-3p, miR-223-5p occur in NIU, JIA, and other autoimmune diseases that often are associated with uveitis (rheumatoid arthritis (RA) [95,96], ankylosing spondylitis (AS) [97], ulcerative colitis (UC) [98], lupus erythematosus (LE) [99]). There are no miRNAs specific only for uveitis or common only for uveitis and juvenile idiopathic arthritis that are not detected in other autoimmune disorders. Created with BioRender.com

**Table 4.**
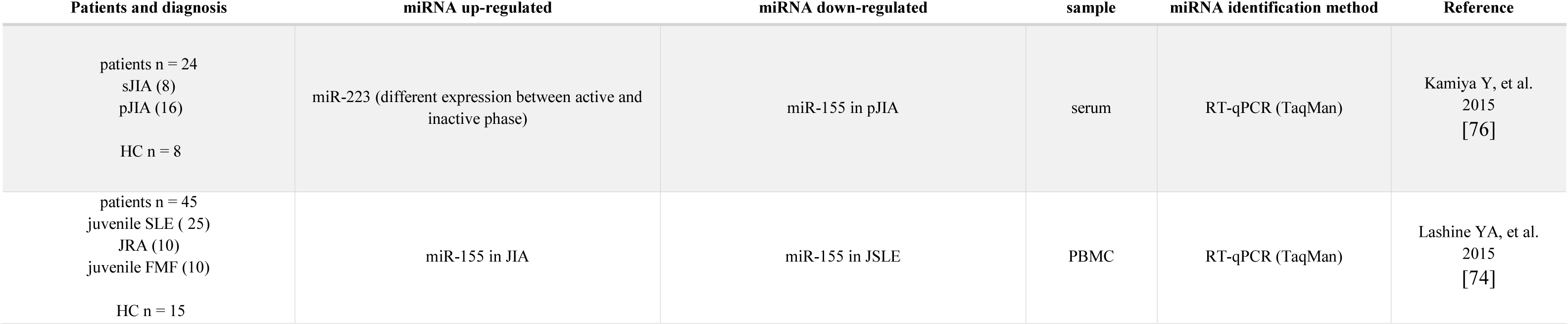
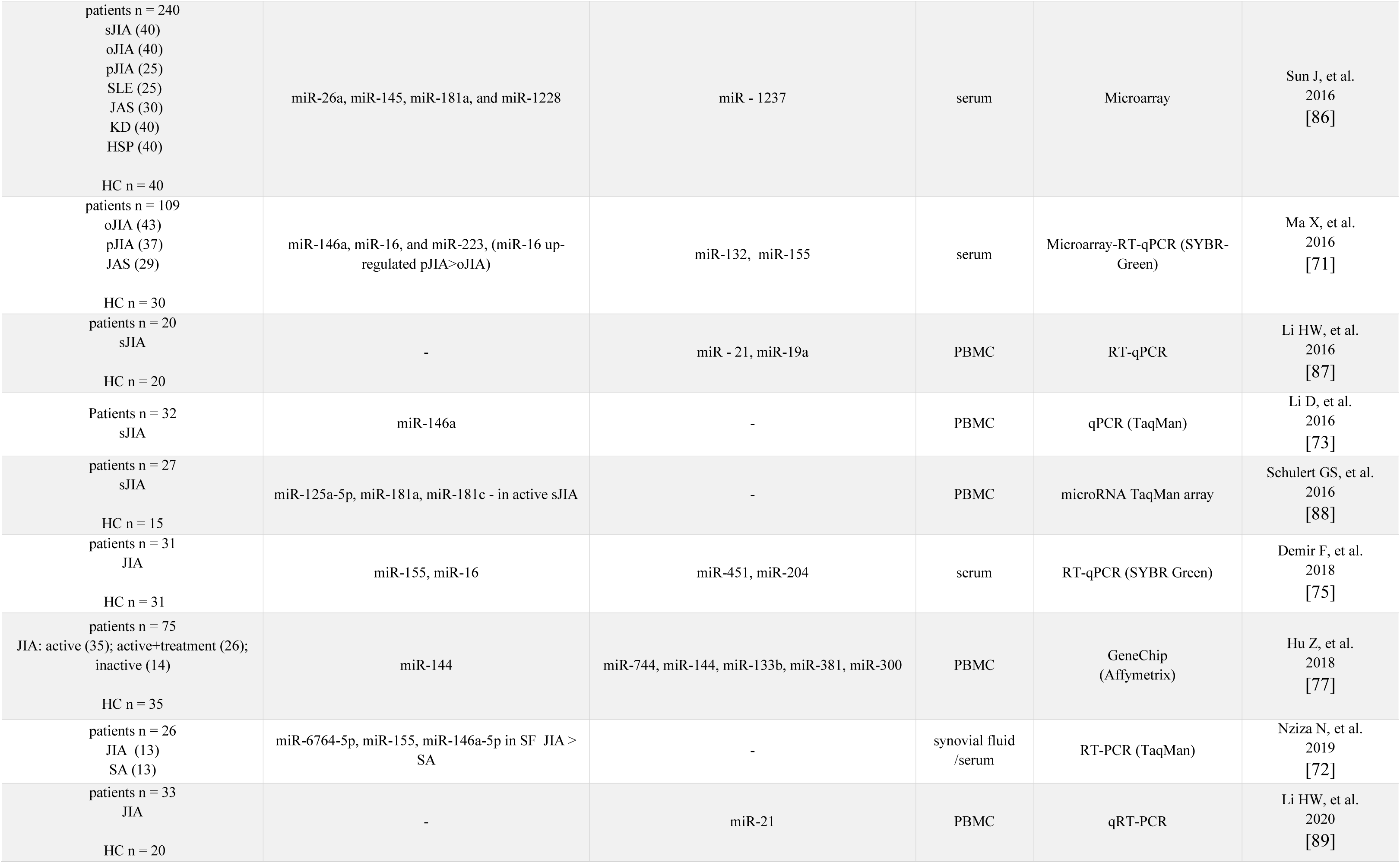
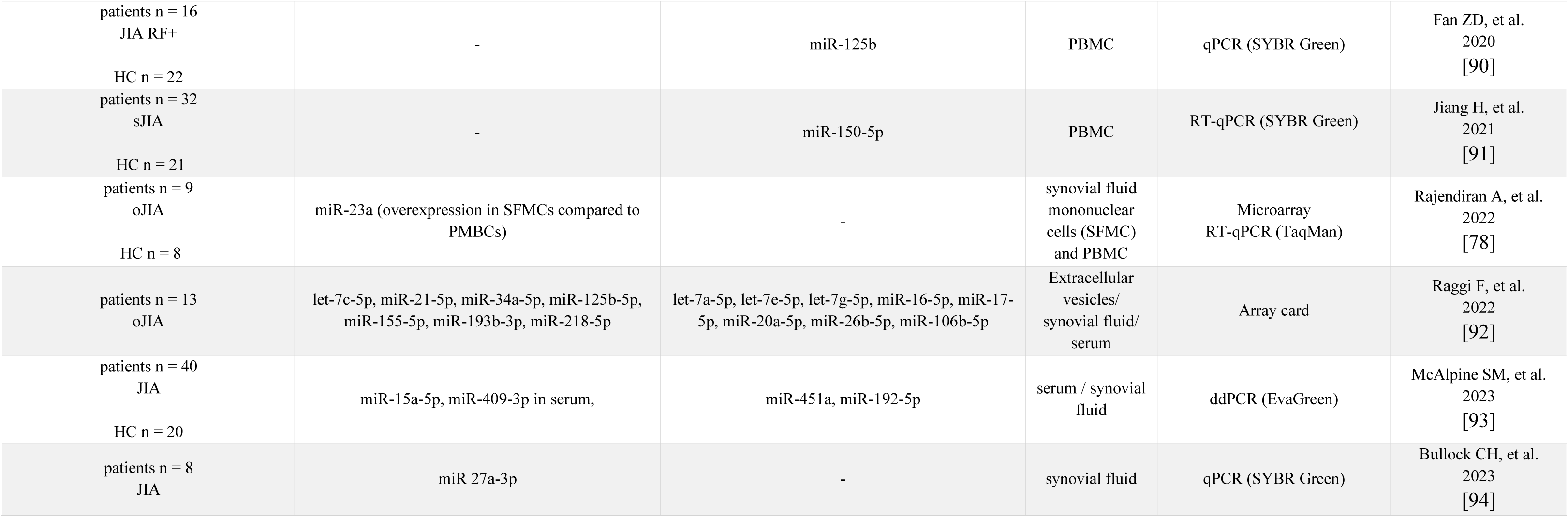
miRNA in juvenile idiopathic arthritis. HC: healthy controls JIA: juvenile idiopathic arthritis; sJIA: systemic JIA; pJIA: polyarticular JIA; oJIA: oligoarticular JIA; JAS: juvenile ankylosing spondylitis; SLE: systemic lupus erythematosus; KD: Kawasaki disease; HSP: Henoch-Schönlein purpura; SA: septic arthritis, PBMC: peripheral blood mononuclear cells, RF: Rheumatoid Factor.

### 3.3 Analysis of miRNAs targets and signaling pathways

To recognize a regulatory function of analyzed miRNAs in non-infectious uveitis pathology, we performed *in silico* target annotation analysis between 10 miRNAs found in patients with uveitis and 247 uveitis associated genes received from DisGeNET 7.0 database (Concept Unique Identifier “C0042164” was queried) likewise 9 miRNAs found in patients with juvenile idiopathic arthritis-associated uveitis and 450 juvenile arthritis associated genes received from DisGeNET 7.0 database (Concept Unique Identifier “C3495559” was queried). Target annotation analysis revealed 264 validated miRNA-gene pairs for uveitis, including 142 unique genes. (**Supplementary Table 4**). Of these, 74 genes were regulated by more than one of the identified miRNAs, with one gene being target of six of the miRNAs (*VEGFA*), 2 genes being targets of fifth miRNAs (*VCAN*, *IL6ST*), 10 genes being targets of fourth miRNAs, 16 genes being targets of three miRNAs, and 45 genes being targets of two miRNAs. In case of idiopathic arthritis-associated uveitis, target annotation analysis revealed 648 validated miRNA-gene pairs, including 299 unique genes. (**Supplementary Table 5**). Of these, 193 genes were regulated by more than one of the identified miRNAs, with six genes being targets of six of the miRNAs (*VEGFA, IL6, IL6ST, JUN, STAT1, STAT3*), 12 genes being targets of fifth miRNAs, 20 genes being targets of fourth miRNAs, 56 genes being targets of three miRNAs, and 99 genes being targets of two miRNAs. Identified interactions (genes regulated by more than one of the identified miRNAs) were visualized on the regulatory network (**Figure 5**).

**Figure 5.**
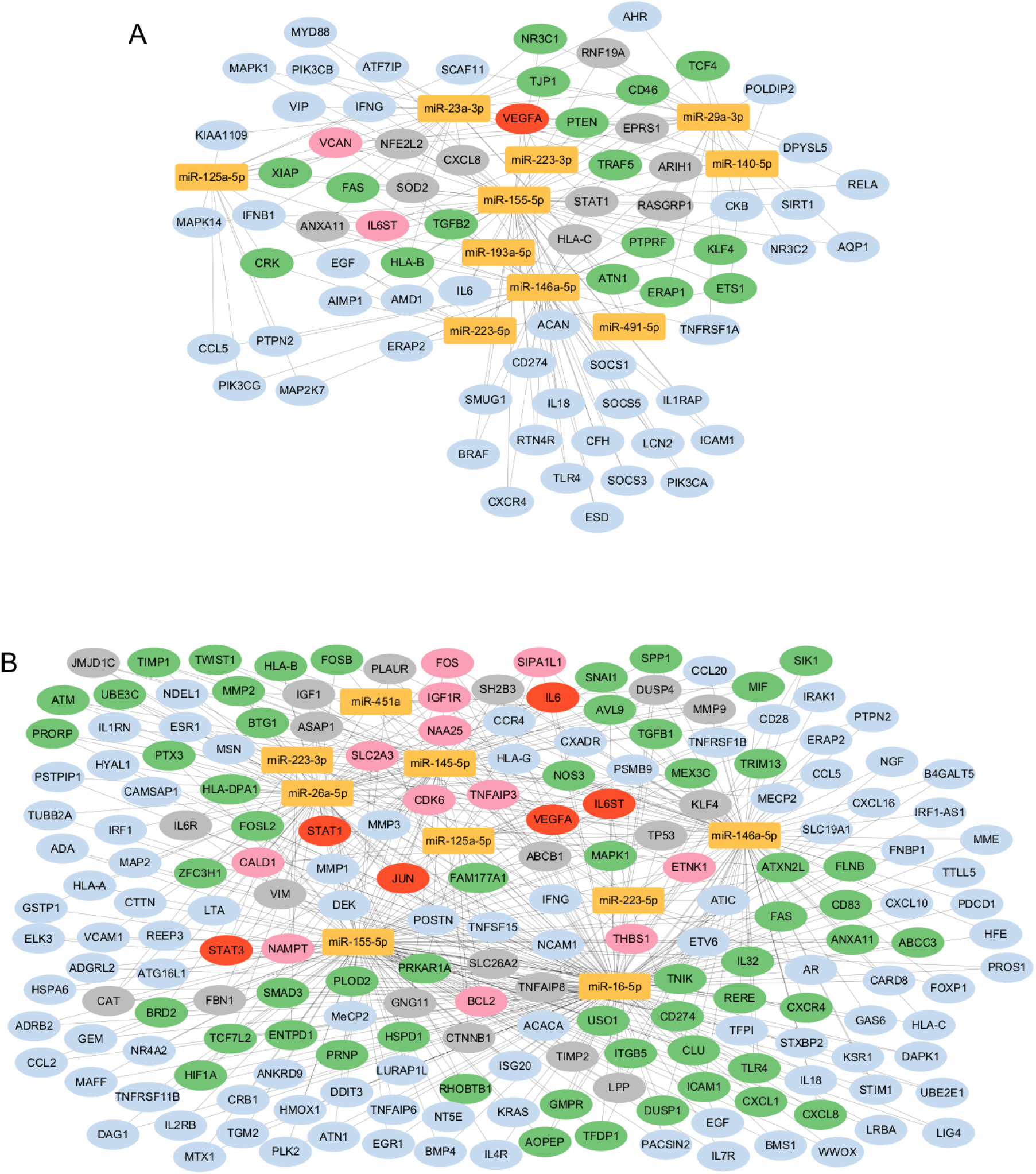
Regulatory network presenting interactions between ten most frequently expressed miRNAs in uveitis. (A) also nine most frequently expressed miRNAs in juvenile idiopathic arthritis-associated uveitis (B) and mRNA targets obtained from DisGeNET 7.0 database as associated with uveitis and juvenile idiopathic arthritis. miRNAs are depicted in yellow circles and the mRNA targets are depicted in blue (targeted by two miRNAs), green (targeted by three miRNAs), grey (targeted by four miRNAs), pink (targeted by fifth miRNAs) and red (targeted by six miRNAs) circles. Interactions were found *in silico* using multiMiR 1.26.0 package in R.

To indicate biological processes in which miRNA-genes are involved, functional analysis was performed for 142 networked genes (uveitis) and 299 networked genes (juvenile idiopathic arthritis-associated uveitis) using the ToppGene Suite website tool (**Supplementary Table 6 and 7**). **Figure 6** presents the top five most enriched terms of GO (Gene Ontology) Biological Processing, GO Molecular Function, GO Cellular Compartment, KEGG, and REACTOME categories.

**Figure 6.**
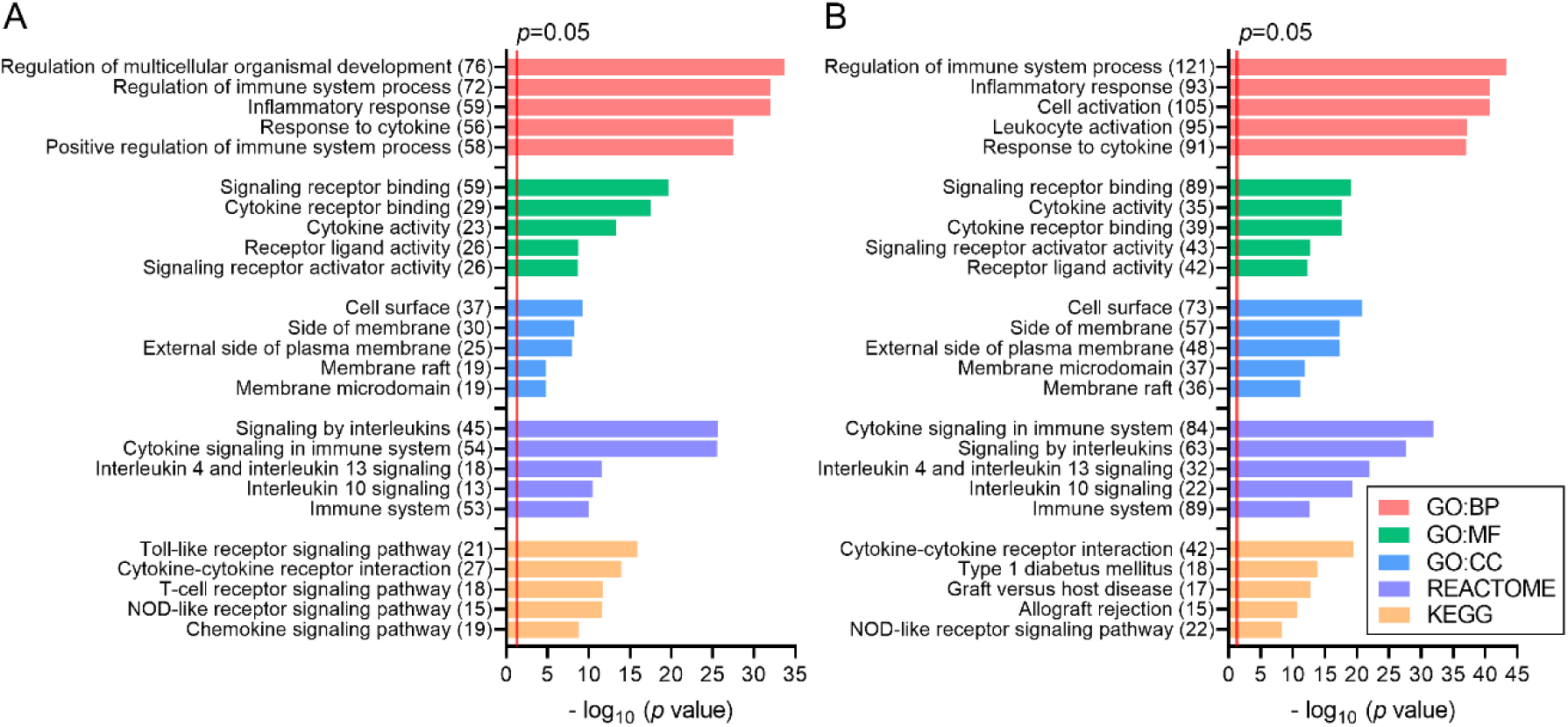
Top five the most enriched terms of GO (Gene Ontology) Biological Processing (GO: BP), GO Molecular Function (GO: MF), GO Cellular Compartment (GO: CC), KEGG (Kyoto Encyclopedia of Genes and Genomes), and REACTOME categories, revealed for uveitis (A) and juvenile idiopathic arthritis-associated uveitis (B) associated genes targeted by miRNAs found in the current study. *p-value* – EASE score for enrichment adjusted by Benjamini correction for multiple hypothesis testing. The number in brackets following the name of terms indicates the number of associated genes.

Genes found as targets of the most frequently expressed miRNAs in both uveitis and juvenile idiopathic arthritis-associated uveitis were associated with the regulation of immune system process and inflammation, response to cytokines, inflammatory response, cell surface, signalling by interleukins, cytokine-cytokine receptor interactions, Toll-like, T-cell, NOD-like receptor, VEGF and JAK/STAT signalling pathways.

## 4. Discussion

Noncoding RNAs regulate multiple cellular processes in every organ in the human body. They promote cellular development and maturation as well as pathological processes. Their involvement in immunological pathways is well established, but their role in childhood autoimmune uveitis is poorly understood and needs to be explored. Non-infectious uveitis in children is commonly idiopathic or associated with JIA. The miRNA environment in this rheumatic disease is somewhat better understood than in uveitis, but to our knowledge, there is no research investigating the role of non-coding RNAs in pediatric idiopathic or autoimmune uveitis or exploring the common background of IU and JIA-AU.

In this study, we analyzed the expression profile of a set of microRNAs in the blood of pediatric uveitis patients and supplemented with data obtained from the meta-analysis. We focused on studies conducted on human samples, mainly because we were looking for miRNA that could be a potential biomarker of idiopathic uveitis. Therefore the sample collection should be as non-invasive, repeatable, and simple as possible. Experimental autoimmune uveitis (EAU) tests are often based on eye tissue samples [24,100]. Although we excluded research on animal models from our meta-analysis, the information on miRNA expression and the course of action that EAU provides are invaluable, especially since the results of miRNA analysis from animal models and human samples are similar and comprise immune-related particles. Several studies have recently reported microRNAs expression profile in adults uveitis patients. However, no one has performed such an analysis on samples from children. Interestingly, our results show similarities with these previous studies but also major differences. Consistent with gene expression studies in adults, a similar miRNA expression pattern in patients with IU may indicate a common molecular basis for this disease with other autoimmune diseases [101,102]. Alterations in miR-30b-5p expression levels are observed in human rheumatoid arthritis and animal models of autoimmune uveitis [103,104]. The overexpression of miR-30b-5p and other ‘inflammatory’ miRNAs in patients P8, P13, and P14 may indicate that uveitis precedes joint symptoms and that these patients may develop JIA or RA in the future. Assessing the accuracy of the assumptions made requires long-term patient follow-up and further research on changes in the expression of selected miRNAs. Downregulation of miR-155-5p in the JIA-AU group (0.30 ± 0.04, *p* = 0.0004) and most patients from the IU group (2.01 ± 0.37, *p* = 0.3863) is consistent with literature findings (**Figure 1**). Similarly, the diverse expression profile of miR-146a-5p is also supported by other research (see **Tables 3** and **4**). miR-146a-5p is involved in several immunological pathways and can be up or downregulated in the serum of patients with autoimmune diseases. Also, the differences between miR-146a-5p expression levels in different papers and our patient group (for JIA-AU patients 0.44 ± 0.04, *p* = 0.0409; for IU patients 1.94 ± 0.18, *p* = 0.2575) (**Figure 1**) may occur because the immune response is an active process and is a constant conversation between immune cells through cytokines and transcription factors creating positive and negative control loops. Therefore, although miR-146a-5p is a key regulator of Th17 development, its expression levels vary at different stages of the immune response [105]. At this point, it may not be a perfect candidate for a molecular marker of the disease. In agreement with the previous publication and the known mechanism of its action, miR-204-5p expression is down-regulated in both JIA-AU (0.12 ± 0.01, *p* < 0.0001) and IU groups (0.15 ± 0.02, *p* < 0.0001) (**Figure 1**) compared to healthy controls [75].

Rosenbaum et al. investigated in the adult population the transcriptional signatures from peripheral blood and identified specific subsets among patients with idiopathic uveitis. Eleven out of 38 patients with idiopathic uveitis had their diagnosis revised based on gene expression profiling. This is especially important when the change of the diagnosis results in a different treatment approach or has different prognostic implications [101]. In juvenile idiopathic arthritis, different subtypes correlate differently with the risk of uveitis and the need for regular ophthalmological checkups. 30% of patients with oligoarticular JIA will present with bilateral, insidious, usually asymptomatic uveitis. In contrast, enthesitis-related arthritis is typically associated with acute anterior uveitis with redness and pain. In both subtypes, the inflammation of the uvea may occur before the joint involvement [16,106]. The “idiopathic” term is usually used in cases where the patient does not fit the diagnostic criteria. We assume, that similar to the adult population, molecular characterization using miRNAs in the pediatric group could provide useful information and allow the reclassification of patients with idiopathic uveitis.

Due to the problem of enrolling a group of patients before any treatment was initiated, we did not correlate the level of inflammatory markers with the selected miRNA expression profiles. Patients with a rheumatological diagnosis had their treatment initiated by rheumatologists, and patients with a diagnosis of idiopathic uveitis also went to the Clinic after initial treatment started by an ophthalmologist at the district clinic. Taking this into account, we carried out an analysis of selected miRNAs in the two patient groups and the healthy control group regardless of the current or past treatment.

Li et al., in gene expression profiling in the adult population, revealed 4 distinct subgroups of uveitis patients. They failed to correlate clinical uveitis diagnoses with this differential gene expression profiling [107]. However, the fact that upregulated genes in one group were more functionally associated with cell proliferation pathways and in the other with cell apoptosis explains why the same therapeutic approach can lead to different outcomes. For example, a monoclonal antibody against interleukin-17 works well in psoriasis but is unsuccessful in rheumatoid arthritis [108,109]. On the other hand, RNA sequencing data show that many forms of uveitis share overlapping mechanisms. Ten upregulated transcripts correlated with immunological pathways and potentially involved in intraocular inflammation were detected in blood samples from patients with different types of uveitis [102]. Identified transcripts include, for example 3 out of 4 myocyte enhancing factor 2 (MEF2) genes. MEF2 protein family is expressed in most tissues, but MEF2B expression is enriched in B cells and other lymphoid tissue. MEF2A, MEF2C, and MEF2D are involved in photoreceptor development and retinal angiogenesis [110,111]. MEF2 family members are targets for miR-223-5p. In chronic myeloid leukemia repression of miR-223 results in the activation of MEF2C and its downregulation negatively correlates with disease risk [112,113]. miR-223 involvement in granulopoiesis and inflammatory signaling cascades is also relevant in JIA and uveitis [66,76].

The eye is an immunologically privileged organ with a unique immunosuppressive environment. For clear vision particularly important is to maintain the immune response under control. The pigmented epithelial (PE) cells of the iris, ciliary body and retina, and corneal endothelial (CE) cells participate in maintaining balance in two ways: firstly, as an anatomical barrier, and secondly, by interacting with active CD4+ cells that have penetrated the blood-ocular barrier via cytokines [114,115]. A disturbance in the balance between Th1, Th2, and Th17 lymphocyte activation and differentiation is the basis of autoimmune uveitis. In addition to the role of these cells, γδ T cells are becoming increasingly prominent in research. They secrete inflammatory cytokines in the initial stages of inflammation, but can also influence the activity of Th17 and escalate the pro-inflammatory effect [116,117].

As shown in **Figure 7**, the involvement of miRNA in immunological regulatory pathways occurs at multiple levels and adds complexity to uveitis pathogenesis. The PE and CE cells act as APCs. Through IL-23 they induce naive CD4+ cells to differentiate into Th17 cells, and downregulation of miR-182-5p additionally enhances this process by the transcriptional initiator TATA-binding protein-associated factor 15 (TAF15) upregulation. Overexpression of miR-182-5p by direct binding with TAF15 mRNA can silence its expression and affect STAT3 levels, which lowers IL-23 receptor (IL-23R) levels as well as retinoid-related orphan receptor-gamma t (RORγt) and granulocyte macrophage-colony stimulating factor (MG-CSF) resulting in depressed Th17 activity and immune reaction [23]. Forkhead box O3 (FOXO3), a direct target of miR-223-3p, is another transcription factor that negatively regulates IL-23R activity. When miR-223-3p levels are elevated, FOXO3 expression is reduced, resulting in increased IL-23R expression and enhanced Th17 cell activity. This creates a feedback loop where IL-23 signaling upregulates miR-223-3p, further suppressing FOXO3 and perpetuating Th17-mediated inflammation [24]. miR-125a and miR-204-5p downregulation can result in prompt immune response. miR-125a promotes T-reg development by suppressing T cell effector factors such as STAT3 and IFN-γ [118,119]. miR-205-5p targets the TGF receptors (TGFβR1 and TGFβR2) and reduce the expression of pro-inflammatory cytokines (IL-1, TNF-α) and matrix metalloproteinases (MMP2, MMP9) [120–122]. An intriguing attribute of miRNAs is that a single miRNA can interact with multiple targets. miR-146a-5p is primarily activated by the nuclear factor kappa-light-chain-enhancer of activated B cells (NF-κB). It acts in a negative loop by targeting TNF receptor-associated factor 6 (TRAF6) and interleukin-1 receptor-associated kinase 1 (IRAK1) and silencing NF-κB. As a result, it inhibits IL-6 and IL-21 signals in CD4+ T cells and hamper their differentiation into Th17 cells [123] [27]. In addition, miR-146a-5p regulates Th17 development *via* IL-2 production and cell death activation [124]. It also interacts with Nucleotide-binding oligomerization domain-containing protein 1 (NOD1) in γδT cells. Nod1 can trigger an inflammatory response that includes the induction of miR-146a. Subsequently, miR-146a directly suppresses NOD1, acting as a cell-intrinsic brake on the ability of γδ27 T cells to produce both IL-17 and IFN-γ, preventing over-activation of the immune response [125,126]. Similarly to miR-146a, miR-155 plays a complex role in regulating the immune response. The expression pattern varies between studies, as shown in **Tables 3 and 4**. It is specific to both tissue type and duration of autoimmune disease [25]. These reflect the fact that miR-155 can manipulate multiple targets in intracellular pathways. In T cells, it interacts with Toll-like receptor (TLR) pathways at different sites, for example by suppressing myeloid differentiation marker 88 (MyD88), TAK-binding protein (TAB), or receptor-interacting protein (RIP). Reducing the level of miR-155 may contribute to the development of EAAU. To add to the complexity, the pro-inflammatory factor STAT3, controlled indirectly by miR-146a, miR-182-5p, and directly by miR-125a, may also act as a regulator of miR-155, creating an axis promoting the expansion of pathogenic Th17 cells in experimental uveitis [28,100,127].

**Figure 7.**
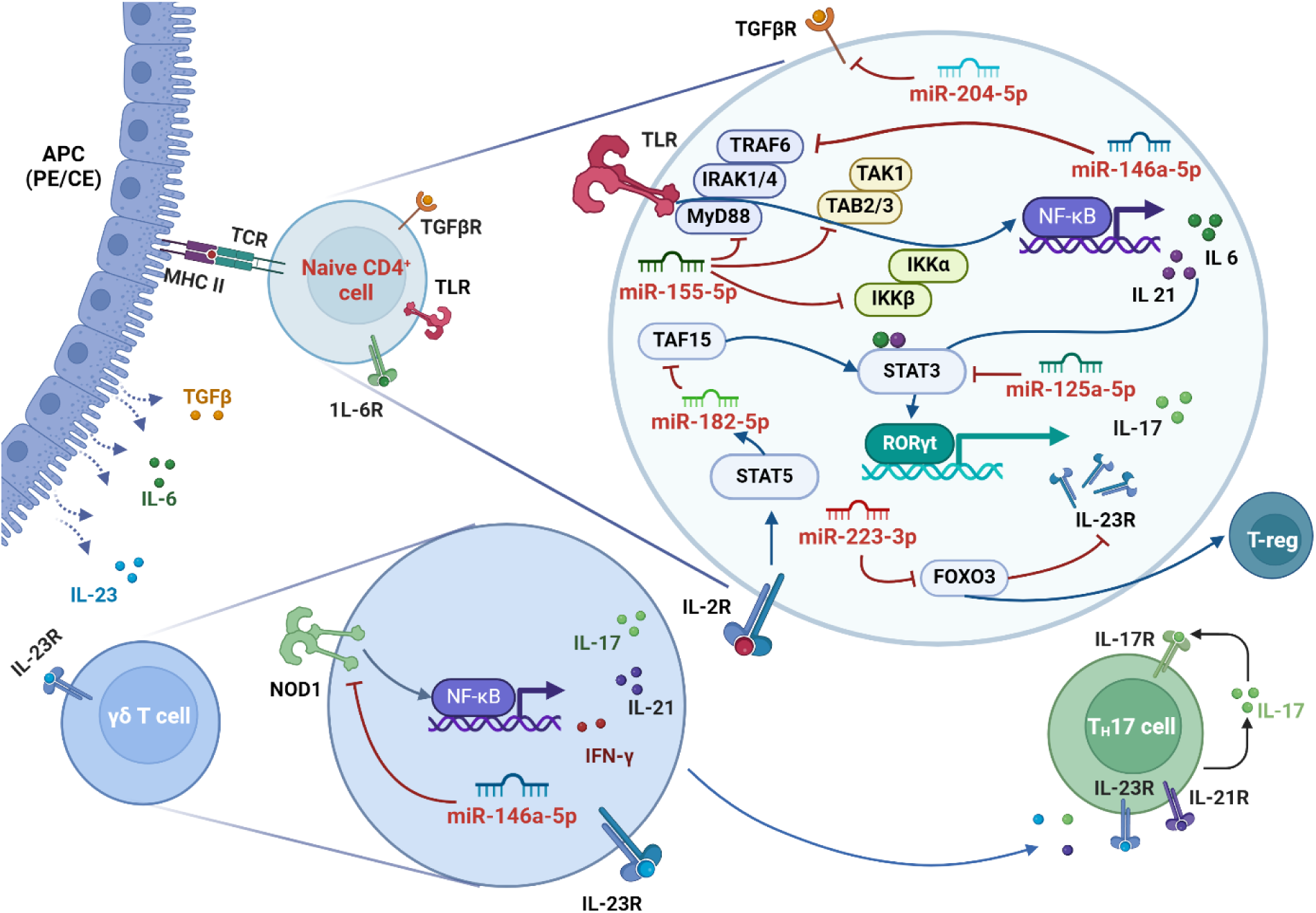
miRNA places of action in T cell maturation after activation by iris pigmented epithelium (PE) or corneal endothelium (CE) acting as antigen-presenting cells (APC). Blue arrows show activation and red flat arrows show inhibition. Detailed description of individual miRNAs – see the text. Created with BioRender.com

Our results have to be interpreted with the following study limitations in mind. The sample collection process was challenging, as it had to be synchronized with lab accessibility and required an additional 10 ml of blood. The main reason patients and their parents/legal guardians did not agree to participate in the study was reluctance to undergo additional blood collection. Also, during the COVID-19 pandemic, the Clinic implemented a restricted admission policy, which complicated patient recruitment and sample collection. As a result, the number of patients included in the study was limited.

Another challenge was the recruitment of a homogenous group of patients. In a tertiary referral center, we observe challenging cases, mainly with chronic or recurrent inflammation resistant to standard treatment and with additional ophthalmological issues. Two of the patients with idiopathic uveitis developed complications, one – secondary glaucoma and one, despite general medications, topical treatment, and extraocular steroid injections – cataract in the left eye. In the JIA group, 6 out of 7 patients were taking a combination of disease-modifying anti-rheumatic drugs (non-biological and biological) at the time of sample collection, and 4 out of 7 had complications such as cataracts or secondary glaucoma.

## 5. Conclusions

Since their discovery, non-coding RNAs such as miRNAs have emerged as master regulators of numerous pathological processes. In the course of idiopathic uveitis, an inflammation of the uvea, many microRNAs show dysregulated expression profile. We confirmed that pediatric idiopathic uveitis shares a common molecular basis with other autoimmune diseases at the non-coding RNA level. The analysis of potential targets also confirmed that the miRNAs studied affect key pathways in T-cell regulation and autoimmunity. Importantly, we showed changes in immune-related miRNA expression levels in patients with a negative rheumatological history and negative ANA antibodies and RF factor. This suggests that alterations at the molecular level are more sensitive and may occur earlier than changes in “classical” antibody levels. These results show that non-coding RNAs are a potential tool to speed up the correct diagnosis and match treatment to an individual patient. Further research is needed to understand the molecular relationships between miRNAs, cytokines, and transcription factors in the complex immune response, especially in an immune-privileged organ like the eye.

## Abbreviations

NIU: non-infectious uveitis
JIA: juvenile idiopathic arthritis
JAS: juvenile ankylosing spondylitis
JSLE: juvenile systemic lupus erythematosus
AIU: autoimmune uveitis
BD: Behcet Disease
RA: rheumatoid arthritis
UC: ulcerative colitis
SLE: systemic lupus erythematosus

## Acknowledgments

We thank all members of the Department of Molecular Neurooncology for their support and discussions. The authors would like to thank Dawid Dorna for his invaluable assistance during the early stages of this project. The authors would like to thank the patients for their collaboration and contribution to this project. Figures were created with the help of GraphPad Prism and BioRender.

## Author contribution

Recruitment and clinical diagnosis of patients, O.W., and A. G.-W, conceptualization, O.W., D.W., A.G.-W., and K.R.; methodology, O.W., D.W., and M.S.; software, O.W., and D.W.; validation, O.W., D.W., and K.R.; formal analysis, O.W, D.W., M.S., and P.G.; investigation, O.W., D.W., M.S., and P.G.; resources, A.G.-W., and K.R.; data curation, O.W., D.W., and K.R.; writing—original draft preparation, O.W., D.W., M.S.; writing—review and editing, O.W., D.W., M.S., A.G.-W., and K.R; visualization, O.W.; supervision, D.W., A.G.-W., and K.R.; project administration, O.W., and D.W.; funding acquisition, A.G.-W., and K.R. All authors read and approved the final version of the manuscript.

## Institutional Review Board Statement

The study was conducted in accordance with the Declaration of Helsinki and approved by the Bioethics Committee of Poznan University of Medical Sciences (951/16 both for patients from the control and case groups).

## Informed Consent Statement

Written prior-informed consent was obtained from the parents/guardians of the patients.

## Data Availability Statement

The data that support the findings of this study are available from the corresponding author upon reasonable request.

## SUPPLEMENTARY MATERIAL

**Supplementary Table 1.** Clinical characteristic of patients.

| **Patient and diagnosis** | **Age at the diagnosis** | **Age at sample collection** | **Sex** | **ANA** | **HLA-B27** | **Additional information /complications** | **Treatment** |
| --- | --- | --- | --- | --- | --- | --- | --- |
| P1 (JIA-AU) | 4 | 10 | M | + | N/A | - | topical medications |
| P2 (JIA-AU) | 3 | 17 | F | + | N/A | AIOL left eye | topical medications<br>MTX, metyloprednizolon, |
| P3 (JIA-AU) | 7 | 13 | M | + | - | AIOL both eyes | topical medications,<br>adalimumab |
| P4 (JIA-AU) | 4 | 10 | F | + | N/A | macular edema | topical medications,<br>MTX |
| P5 (JIA-AU) | 12 | 17 | M | + | N/A | secondary glaucoma | topical medications,<br>MTX, cyclosporin |
| P6 (JIA-AU) | 3 | 17 | F | + | N/A | - | topical medications,<br>MTX, etanercept,<br>adalimumab |
| P7 (JIA-AU) | 12 | 14 | F | + | + | - | topical medications<br>adalimumab, tofacitynib |
| P8 (IU) | 12 | 12 | F | - | - | - | topical medications |
| P9 (IU) | 16 | 17 | M | N/A | N/A | - | topical medications |
| P10 (IU) | 8 | 17 | M | - | - | AIOL left eye | topical medications,<br>prednizon |
| P11 (IU) | 15 | 15 | F | - | - | - | topical medications |
| P12 (IU) | 10 | 13 | M | + | - | - | topical medications |
| P13 (IU) | 12 | 14 | M | - | - | secondary glaucoma | topical medications |
| P14 (IU) | 16 | 17 | F | N/A | N/A | - | topical medications |
| P15 (IU) | 13 | 14 | M | N/A | N/A | - | topical medications |

JIA-AU - juvenile idiopathic arthritis-associated uveitis, IU - Idiopathic uveitis, +/- - positive/negative ANA or HLA-B27 status, MTX – methotrexate, AIOL - artificial intraocular lens, M - male, F - female.

**Supplementary Table 2.**
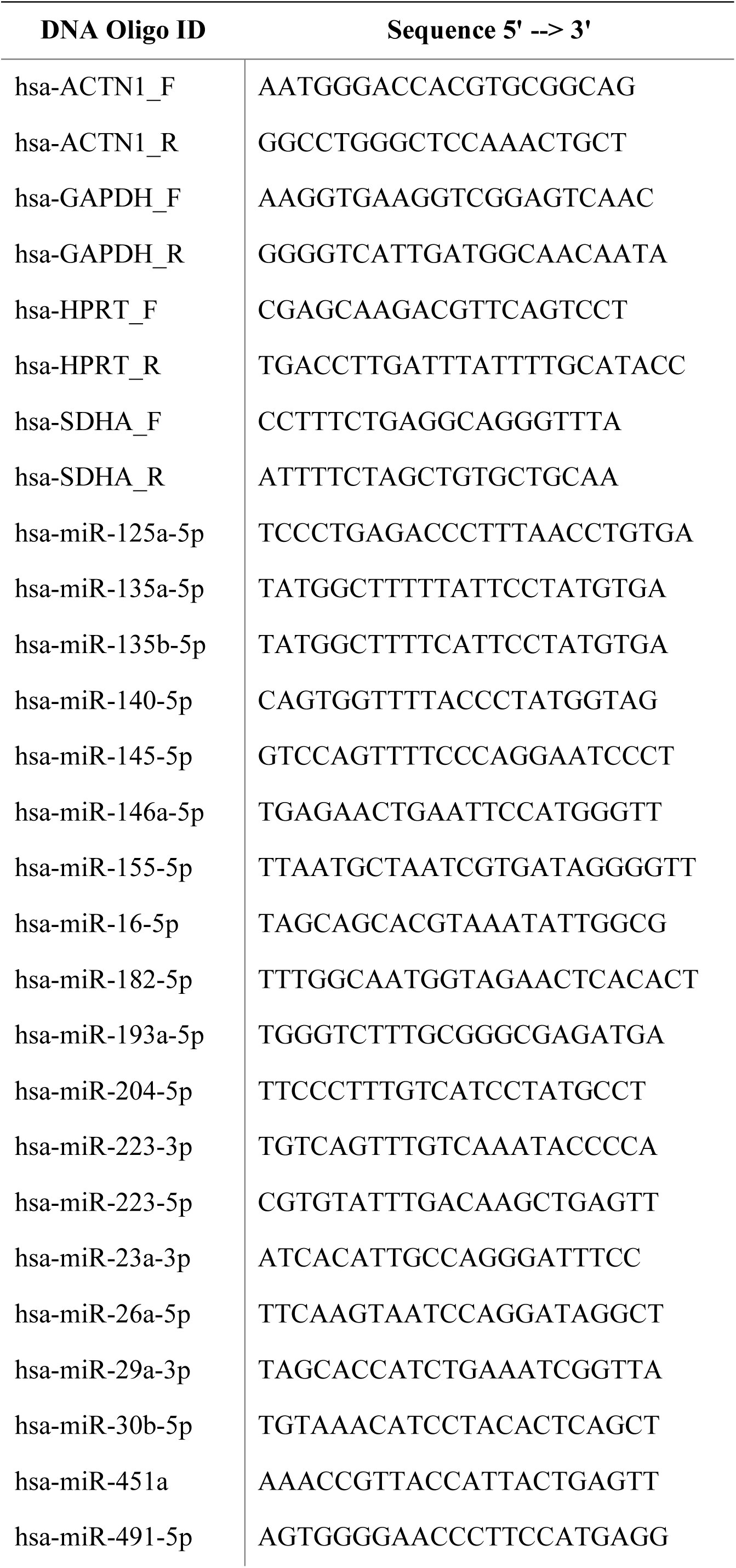
List of primers used in the study.

**Supplementary Table 3.**
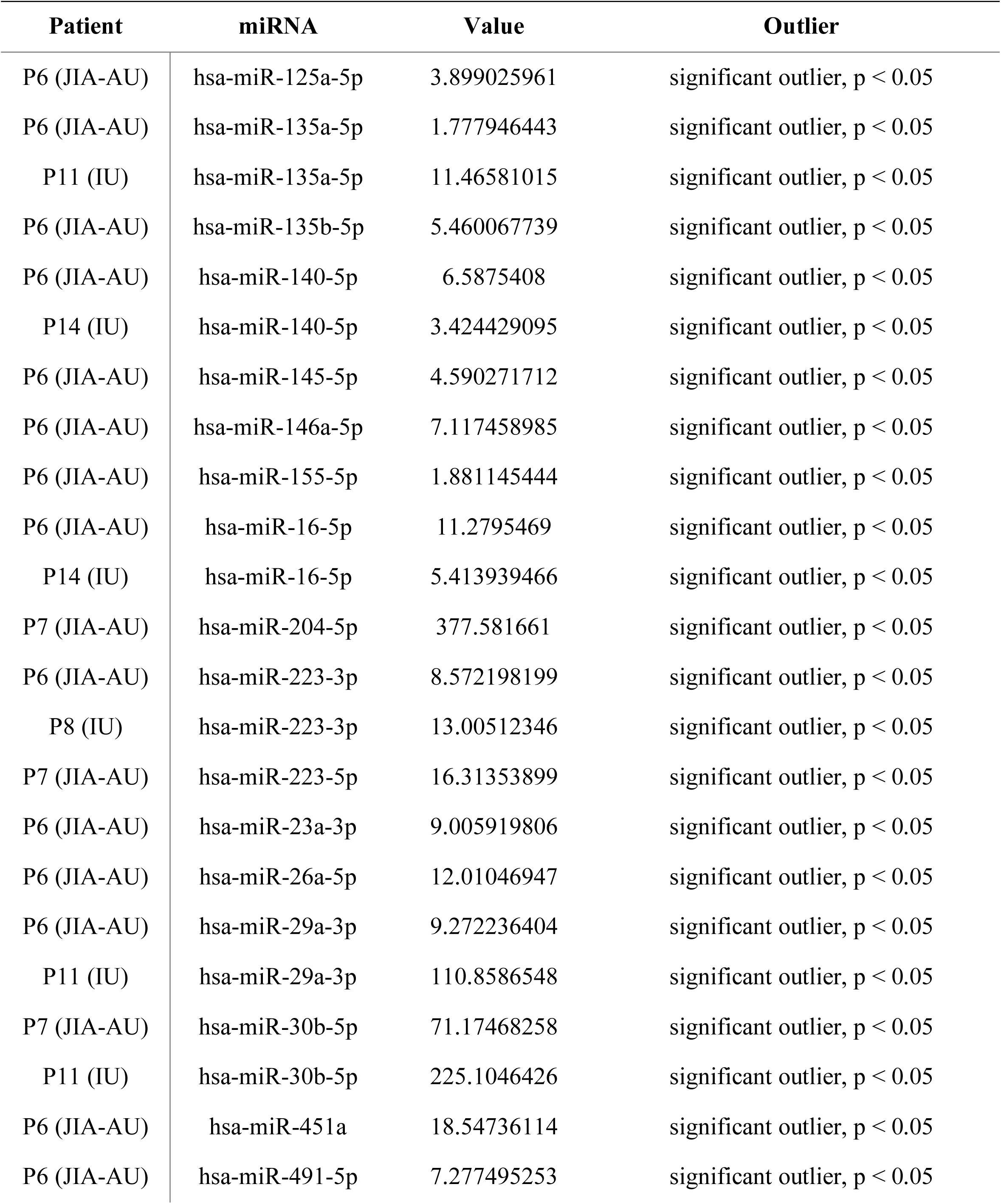
List of significant outliers. Data not shown on Figure 2.

**Supplementary Table 4.**
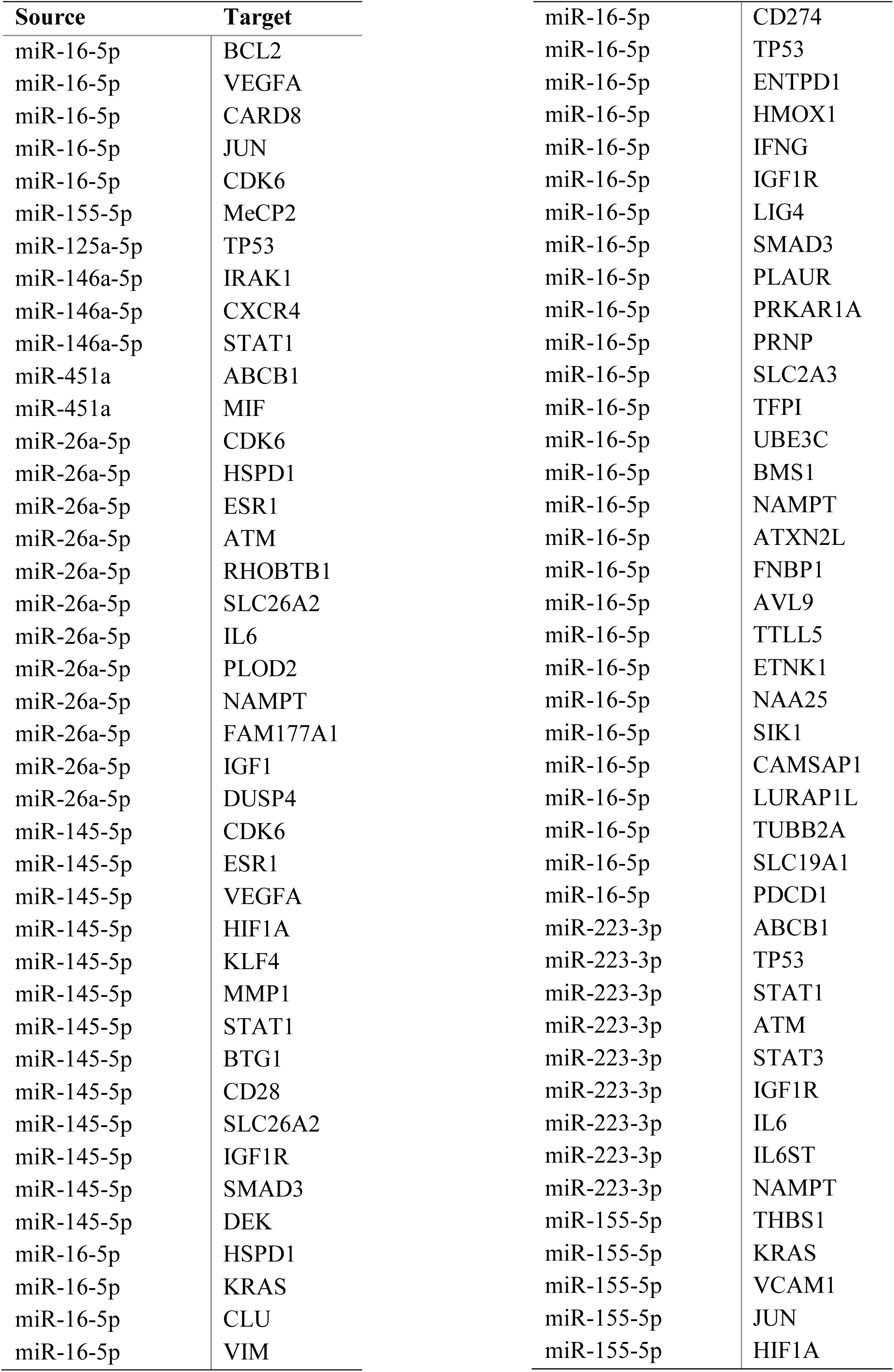
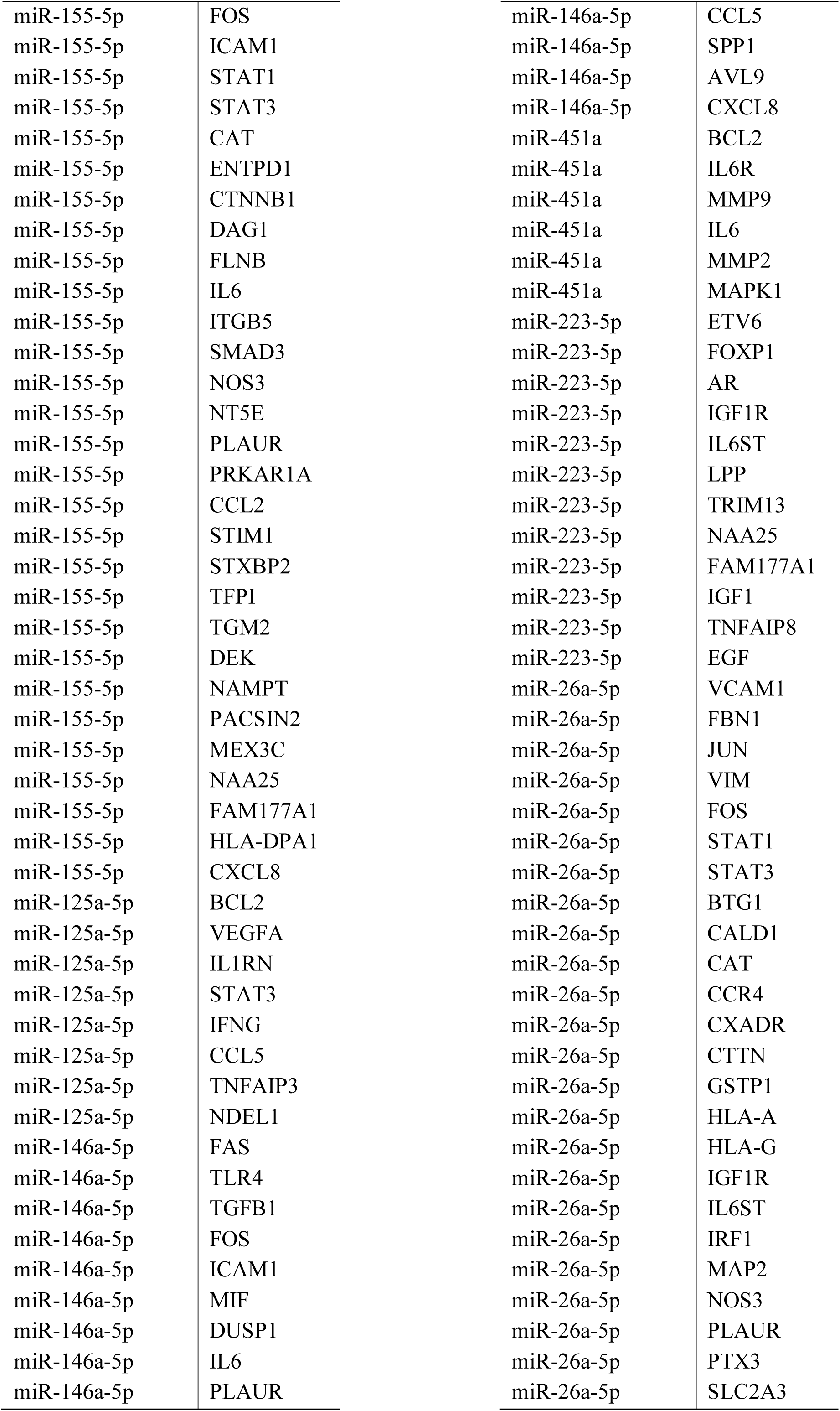
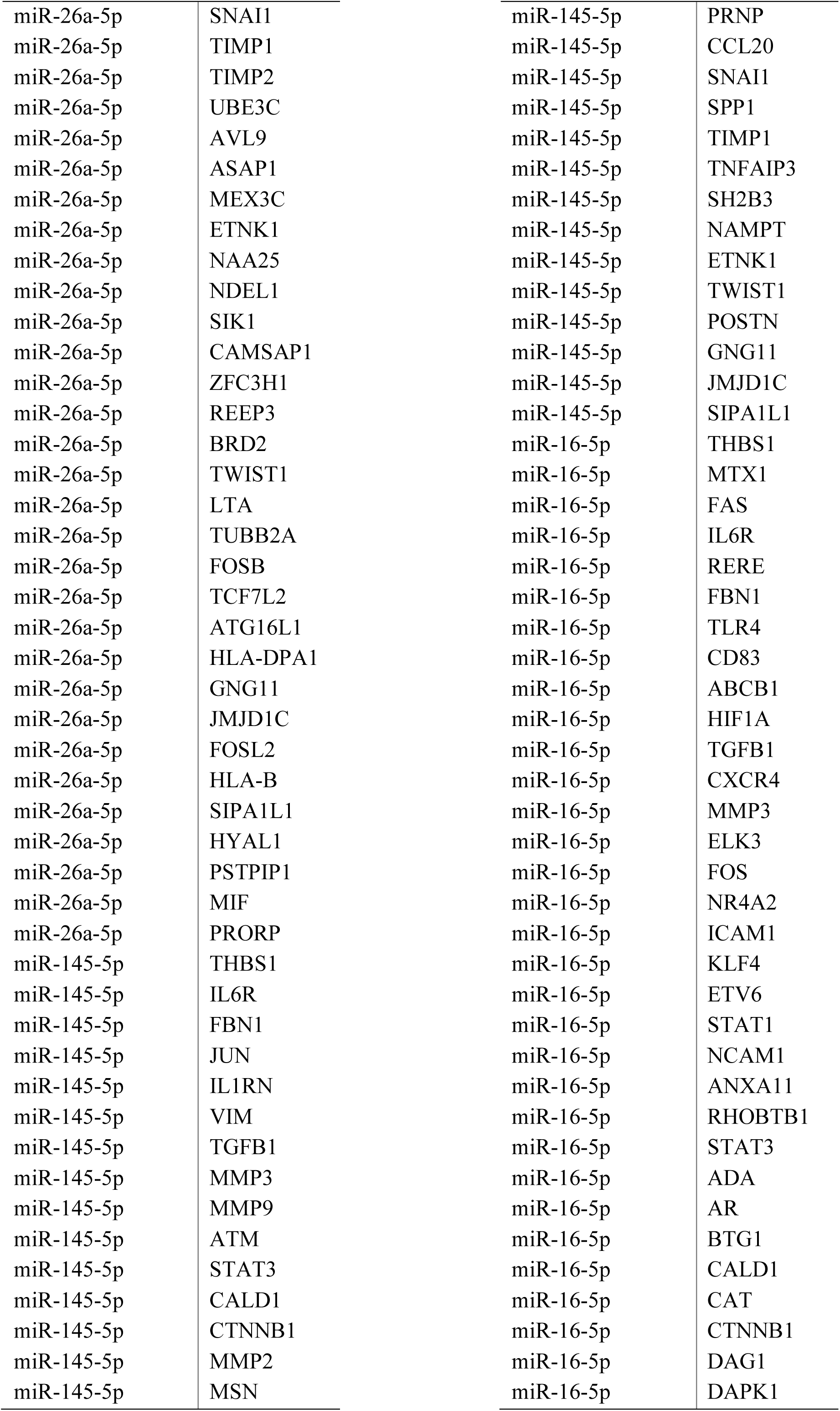
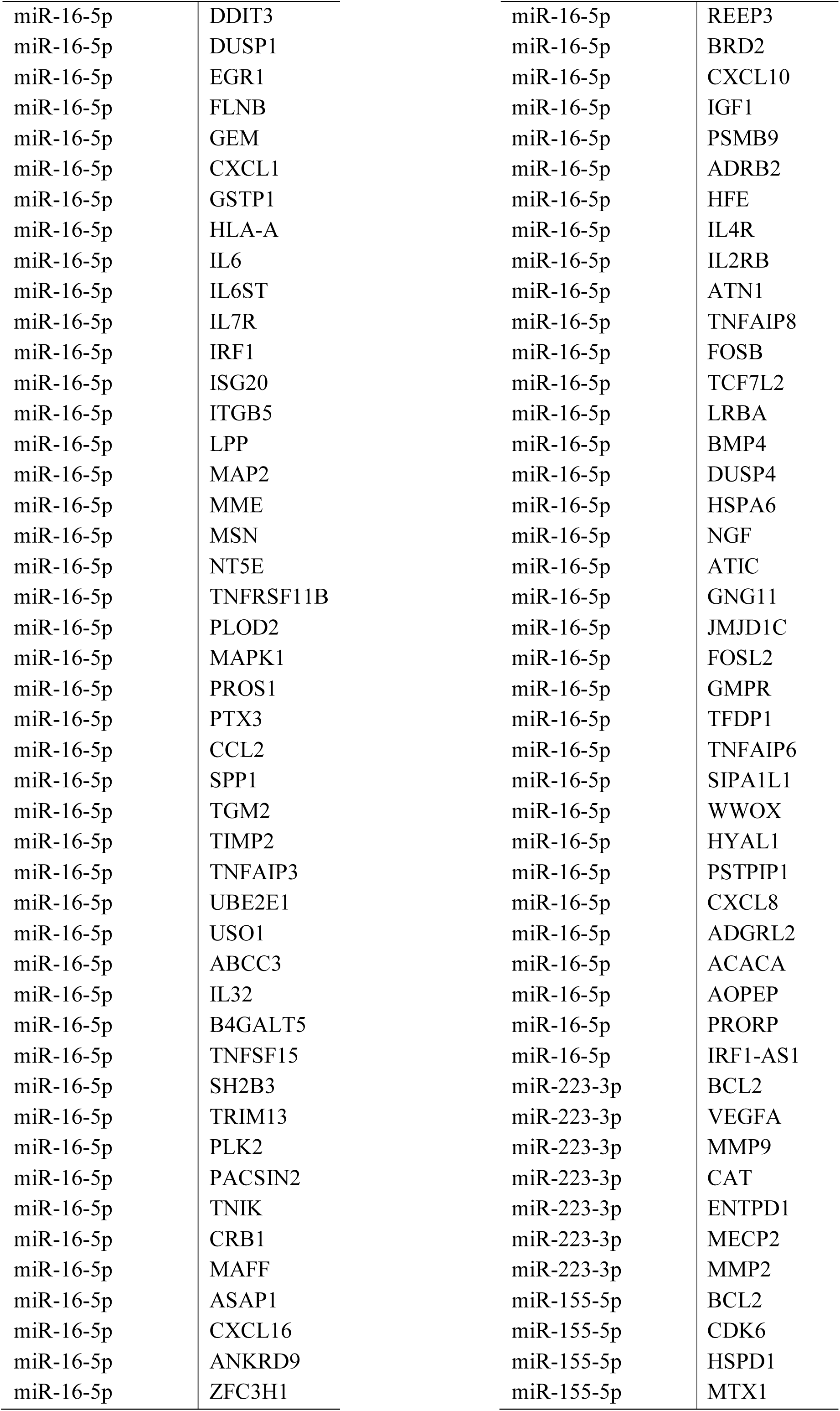
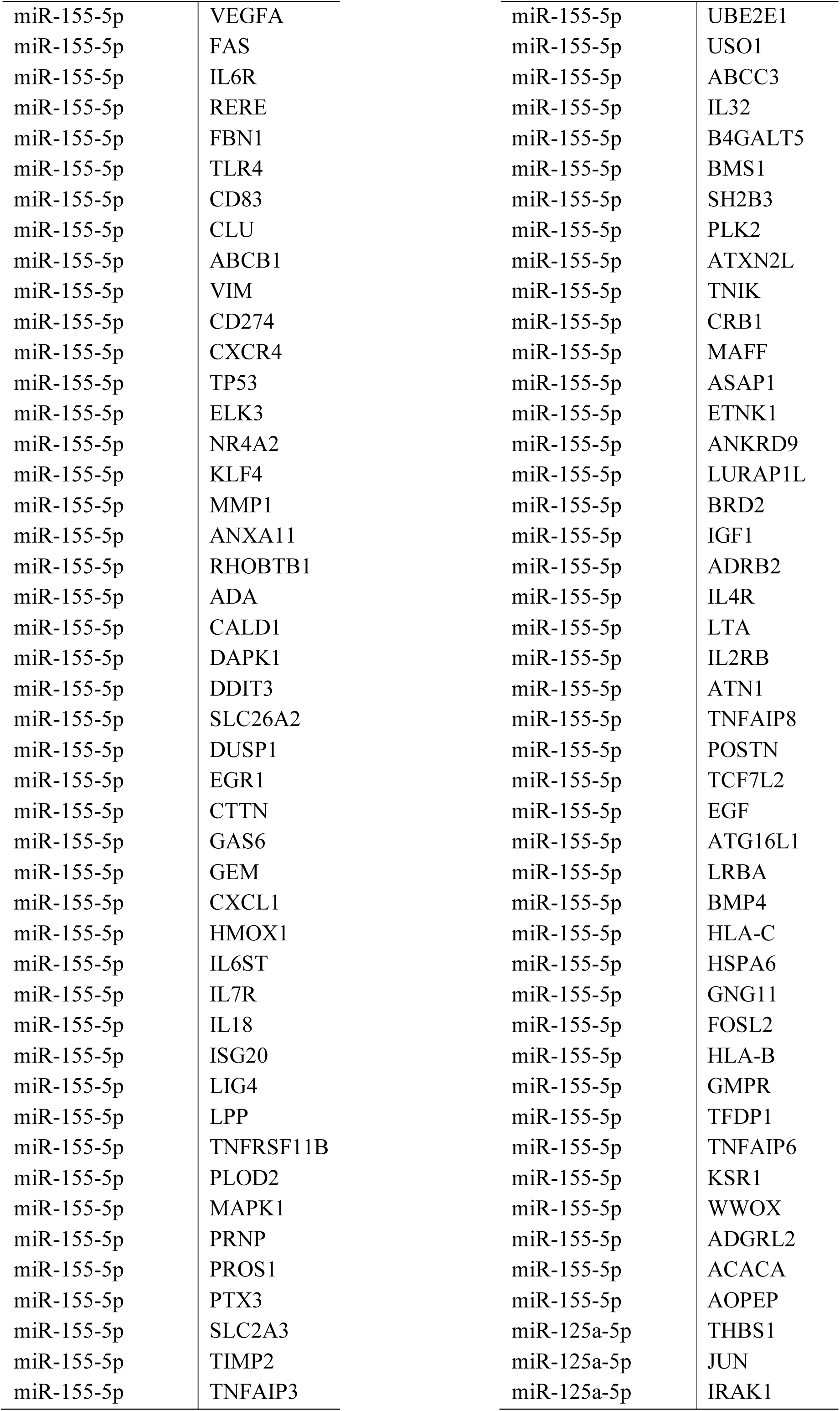
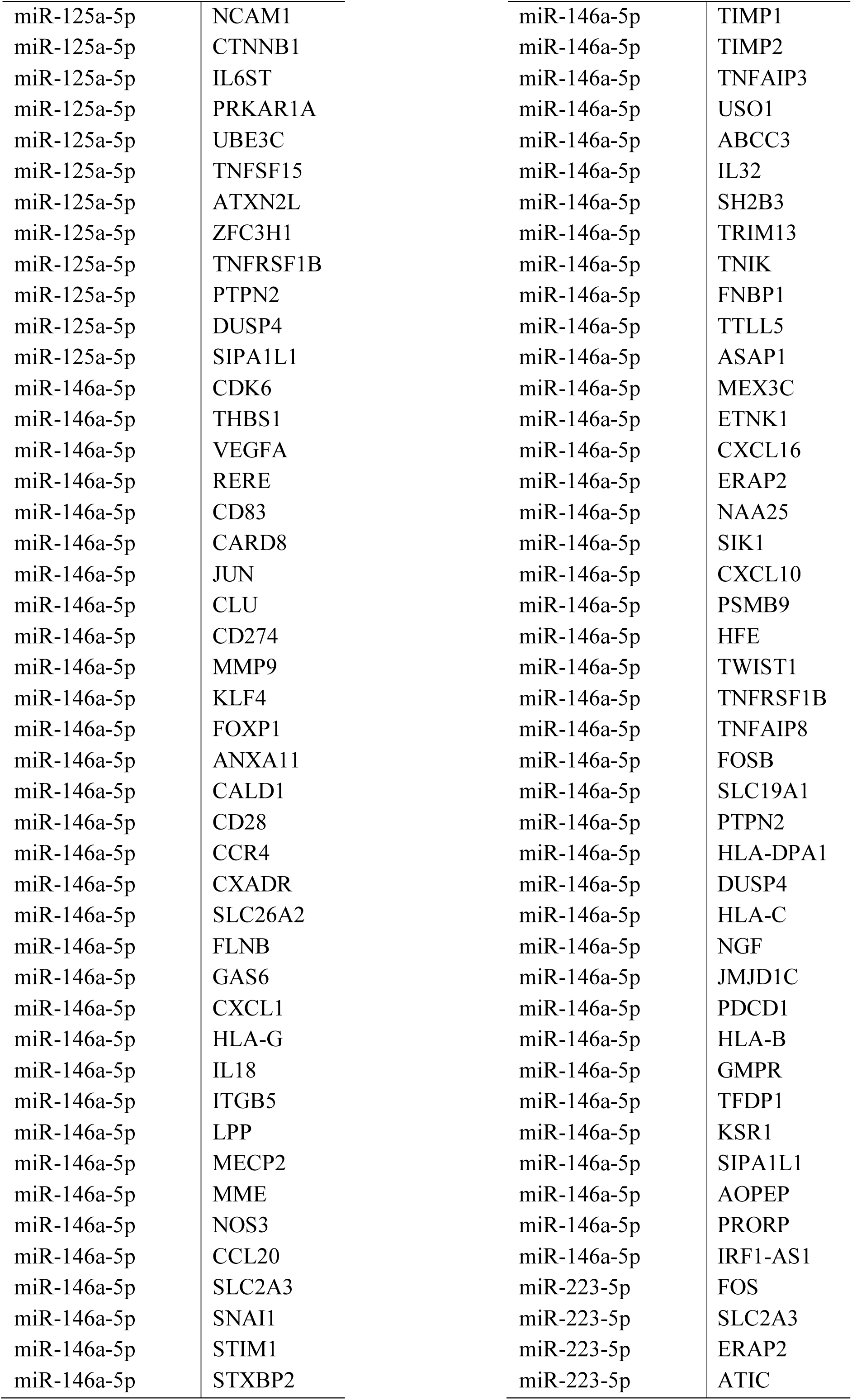
List of genes (193) regulated by more than one of the identified miRNAs in patients juvenile arthritis-associated uveitis (541 miRNA-gene interaction pairs).

| Source | Target |  |  |
| --- | --- | --- | --- |
| miR-16-5p | BCL2 | miR-16-5p | CD274 |
| miR-16-5p | VEGFA | miR-16-5p | TP53 |
| miR-16-5p | CARD8 | miR-16-5p | ENTPD1 |
| miR-16-5p | JUN | miR-16-5p | HMOX1 |
| miR-16-5p | CDK6 | miR-16-5p | IFNG |
| miR-155-5p | MeCP2 | miR-16-5p | IGF1R |
| miR-125a-5p | TP53 | miR-16-5p | LIG4 |
| miR-146a-5p | IRAK1 | miR-16-5p | SMAD3 |
| miR-146a-5p | CXCR4 | miR-16-5p | PLAUR |
| miR-146a-5p | STAT1 | miR-16-5p | PRKAR1A |
| miR-451a | ABCB1 | miR-16-5p | PRNP |
| miR-451a | MIF | miR-16-5p | SLC2A3 |
| miR-26a-5p | CDK6 | miR-16-5p | TFPI |
| miR-26a-5p | HSPD1 | miR-16-5p | UBE3C |
| miR-26a-5p | ESR1 | miR-16-5p | BMS1 |
| miR-26a-5p | ATM | miR-16-5p | NAMPT |
| miR-26a-5p | RHOBTB1 | miR-16-5p | ATXN2L |
| miR-26a-5p | SLC26A2 | miR-16-5p | FNBP1 |
| miR-26a-5p | IL6 | miR-16-5p | AVL9 |
| miR-26a-5p | PLOD2 | miR-16-5p | TTLL5 |
| miR-26a-5p | NAMPT | miR-16-5p | ETNK1 |
| miR-26a-5p | FAM177A1 | miR-16-5p | NAA25 |
| miR-26a-5p | IGF1 | miR-16-5p | SIK1 |
| miR-26a-5p | DUSP4 | miR-16-5p | CAMSAP1 |
| miR-145-5p | CDK6 | miR-16-5p | LURAP1L |
| miR-145-5p | ESR1 | miR-16-5p | TUBB2A |
| miR-145-5p | VEGFA | miR-16-5p | SLC19A1 |
| miR-145-5p | HIF1A | miR-16-5p | PDCD1 |
| miR-145-5p | KLF4 | miR-223-3p | ABCB1 |
| miR-145-5p | MMP1 | miR-223-3p | TP53 |
| miR-145-5p | STAT1 | miR-223-3p | STAT1 |
| miR-145-5p | BTG1 | miR-223-3p | ATM |
| miR-145-5p | CD28 | miR-223-3p | STAT3 |
| miR-145-5p | SLC26A2 | miR-223-3p | IGF1R |
| miR-145-5p | IGF1R | miR-223-3p | IL6 |
| miR-145-5p | SMAD3 | miR-223-3p | IL6ST |
| miR-145-5p | DEK | miR-223-3p | NAMPT |
| miR-16-5p | HSPD1 | miR-155-5p | THBS1 |
| miR-16-5p | KRAS | miR-155-5p | KRAS |
| miR-16-5p | CLU | miR-155-5p | VCAM1 |
| miR-16-5p | VIM | miR-155-5p | JUN |
|  |  | miR-155-5p | HIF1A |

|  |  |
| --- | --- |
| miR-155-5p | FOS |
| miR-155-5p | ICAM1 |
| miR-155-5p | STAT1 |
| miR-155-5p | STAT3 |
| miR-155-5p | CAT |
| miR-155-5p | ENTPD1 |
| miR-155-5p | CTNNB1 |
| miR-155-5p | DAG1 |
| miR-155-5p | FLNB |
| miR-155-5p | IL6 |
| miR-155-5p | ITGB5 |
| miR-155-5p | SMAD3 |
| miR-155-5p | NOS3 |
| miR-155-5p | NT5E |
| miR-155-5p | PLAUR |
| miR-155-5p | PRKAR1A |
| miR-155-5p | CCL2 |
| miR-155-5p | STIM1 |
| miR-155-5p | STXBP2 |
| miR-155-5p | TFPI |
| miR-155-5p | TGM2 |
| miR-155-5p | DEK |
| miR-155-5p | NAMPT |
| miR-155-5p | PACSIN2 |
| miR-155-5p | MEX3C |
| miR-155-5p | NAA25 |
| miR-155-5p | FAM177A1 |
| miR-155-5p | HLA-DPA1 |
| miR-155-5p | CXCL8 |
| miR-125a-5p | BCL2 |
| miR-125a-5p | VEGFA |
| miR-125a-5p | IL1RN |
| miR-125a-5p | STAT3 |
| miR-125a-5p | IFNG |
| miR-125a-5p | CCL5 |
| miR-125a-5p | TNFAIP3 |
| miR-125a-5p | NDEL1 |
| miR-146a-5p | FAS |
| miR-146a-5p | TLR4 |
| miR-146a-5p | TGFB1 |
| miR-146a-5p | FOS |
| miR-146a-5p | ICAM1 |
| miR-146a-5p | MIF |
| miR-146a-5p | DUSP1 |
| miR-146a-5p | IL6 |
| miR-146a-5p | PLAUR |
| miR-146a-5p | CCL5 |
| miR-146a-5p | SPP1 |
| miR-146a-5p | AVL9 |
| miR-146a-5p | CXCL8 |
| miR-451a | BCL2 |
| miR-451a | IL6R |
| miR-451a | MMP9 |
| miR-451a | IL6 |
| miR-451a | MMP2 |
| miR-451a | MAPK1 |
| miR-223-5p | ETV6 |
| miR-223-5p | FOXP1 |
| miR-223-5p | AR |
| miR-223-5p | IGF1R |
| miR-223-5p | IL6ST |
| miR-223-5p | LPP |
| miR-223-5p | TRIM13 |
| miR-223-5p | NAA25 |
| miR-223-5p | FAM177A1 |
| miR-223-5p | IGF1 |
| miR-223-5p | TNFAIP8 |
| miR-223-5p | EGF |
| miR-26a-5p | VCAM1 |
| miR-26a-5p | FBN1 |
| miR-26a-5p | JUN |
| miR-26a-5p | VIM |
| miR-26a-5p | FOS |
| miR-26a-5p | STAT1 |
| miR-26a-5p | STAT3 |
| miR-26a-5p | BTG1 |
| miR-26a-5p | CALD1 |
| miR-26a-5p | CAT |
| miR-26a-5p | CCR4 |
| miR-26a-5p | CXADR |
| miR-26a-5p | CTTN |
| miR-26a-5p | GSTP1 |
| miR-26a-5p | HLA-A |
| miR-26a-5p | HLA-G |
| miR-26a-5p | IGF1R |
| miR-26a-5p | IL6ST |
| miR-26a-5p | IRF1 |
| miR-26a-5p | MAP2 |
| miR-26a-5p | NOS3 |
| miR-26a-5p | PLAUR |
| miR-26a-5p | PTX3 |
| miR-26a-5p | SLC2A3 |
| miR-26a-5p | SNAI1 |
| miR-26a-5p | TIMP1 |
| miR-26a-5p | TIMP2 |
| miR-26a-5p | UBE3C |
| miR-26a-5p | AVL9 |
| miR-26a-5p | ASAP1 |
| miR-26a-5p | MEX3C |
| miR-26a-5p | ETNK1 |
| miR-26a-5p | NAA25 |
| miR-26a-5p | NDEL1 |
| miR-26a-5p | SIK1 |
| miR-26a-5p | CAMSAP1 |
| miR-26a-5p | ZFC3H1 |
| miR-26a-5p | REEP3 |
| miR-26a-5p | BRD2 |
| miR-26a-5p | TWIST1 |
| miR-26a-5p | LTA |
| miR-26a-5p | TUBB2A |
| miR-26a-5p | FOSB |
| miR-26a-5p | TCF7L2 |
| miR-26a-5p | ATG16L1 |
| miR-26a-5p | HLA-DPA1 |
| miR-26a-5p | GNG11 |
| miR-26a-5p | JMJD1C |
| miR-26a-5p | FOSL2 |
| miR-26a-5p | HLA-B |
| miR-26a-5p | SIPA1L1 |
| miR-26a-5p | HYAL1 |
| miR-26a-5p | PSTPIP1 |
| miR-26a-5p | MIF |
| miR-26a-5p | PRORP |
| miR-145-5p | THBS1 |
| miR-145-5p | IL6R |
| miR-145-5p | FBN1 |
| miR-145-5p | JUN |
| miR-145-5p | IL1RN |
| miR-145-5p | VIM |
| miR-145-5p | TGFB1 |
| miR-145-5p | MMP3 |
| miR-145-5p | MMP9 |
| miR-145-5p | ATM |
| miR-145-5p | STAT3 |
| miR-145-5p | CALD1 |
| miR-145-5p | CTNNB1 |
| miR-145-5p | MMP2 |
| miR-145-5p | MSN |
| miR-145-5p | PRNP |
| miR-145-5p | CCL20 |
| miR-145-5p | SNAI1 |
| miR-145-5p | SPP1 |
| miR-145-5p | TIMP1 |
| miR-145-5p | TNFAIP3 |
| miR-145-5p | SH2B3 |
| miR-145-5p | NAMPT |
| miR-145-5p | ETNK1 |
| miR-145-5p | TWIST1 |
| miR-145-5p | POSTN |
| miR-145-5p | GNG11 |
| miR-145-5p | JMJD1C |
| miR-145-5p | SIPA1L1 |
| miR-16-5p | THBS1 |
| miR-16-5p | MTX1 |
| miR-16-5p | FAS |
| miR-16-5p | IL6R |
| miR-16-5p | RERE |
| miR-16-5p | FBN1 |
| miR-16-5p | TLR4 |
| miR-16-5p | CD83 |
| miR-16-5p | ABCB1 |
| miR-16-5p | HIF1A |
| miR-16-5p | TGFB1 |
| miR-16-5p | CXCR4 |
| miR-16-5p | MMP3 |
| miR-16-5p | ELK3 |
| miR-16-5p | FOS |
| miR-16-5p | NR4A2 |
| miR-16-5p | ICAM1 |
| miR-16-5p | KLF4 |
| miR-16-5p | ETV6 |
| miR-16-5p | STAT1 |
| miR-16-5p | NCAM1 |
| miR-16-5p | ANXA11 |
| miR-16-5p | RHOBTB1 |
| miR-16-5p | STAT3 |
| miR-16-5p | ADA |
| miR-16-5p | AR |
| miR-16-5p | BTG1 |
| miR-16-5p | CALD1 |
| miR-16-5p | CAT |
| miR-16-5p | CTNNB1 |
| miR-16-5p | DAG1 |
| miR-16-5p | DAPK1 |
| miR-16-5p | DDIT3 |
| miR-16-5p | DUSP1 |
| miR-16-5p | EGR1 |
| miR-16-5p | FLNB |
| miR-16-5p | GEM |
| miR-16-5p | CXCL1 |
| miR-16-5p | GSTP1 |
| miR-16-5p | HLA-A |
| miR-16-5p | IL6 |
| miR-16-5p | IL6ST |
| miR-16-5p | IL7R |
| miR-16-5p | IRF1 |
| miR-16-5p | ISG20 |
| miR-16-5p | ITGB5 |
| miR-16-5p | LPP |
| miR-16-5p | MAP2 |
| miR-16-5p | MME |
| miR-16-5p | MSN |
| miR-16-5p | NT5E |
| miR-16-5p | TNFRSF11B |
| miR-16-5p | PLOD2 |
| miR-16-5p | MAPK1 |
| miR-16-5p | PROS1 |
| miR-16-5p | PTX3 |
| miR-16-5p | CCL2 |
| miR-16-5p | SPP1 |
| miR-16-5p | TGM2 |
| miR-16-5p | TIMP2 |
| miR-16-5p | TNFAIP3 |
| miR-16-5p | UBE2E1 |
| miR-16-5p | USO1 |
| miR-16-5p | ABCC3 |
| miR-16-5p | IL32 |
| miR-16-5p | B4GALT5 |
| miR-16-5p | TNFSF15 |
| miR-16-5p | SH2B3 |
| miR-16-5p | TRIM13 |
| miR-16-5p | PLK2 |
| miR-16-5p | PACSIN2 |
| miR-16-5p | TNIK |
| miR-16-5p | CRB1 |
| miR-16-5p | MAFF |
| miR-16-5p | ASAP1 |
| miR-16-5p | CXCL16 |
| miR-16-5p | ANKRD9 |
| miR-16-5p | ZFC3H1 |
| miR-16-5p | REEP3 |
| miR-16-5p | BRD2 |
| miR-16-5p | CXCL10 |
| miR-16-5p | IGF1 |
| miR-16-5p | PSMB9 |
| miR-16-5p | ADRB2 |
| miR-16-5p | HFE |
| miR-16-5p | IL4R |
| miR-16-5p | IL2RB |
| miR-16-5p | ATN1 |
| miR-16-5p | TNFAIP8 |
| miR-16-5p | FOSB |
| miR-16-5p | TCF7L2 |
| miR-16-5p | LRBA |
| miR-16-5p | BMP4 |
| miR-16-5p | DUSP4 |
| miR-16-5p | HSPA6 |
| miR-16-5p | NGF |
| miR-16-5p | ATIC |
| miR-16-5p | GNG11 |
| miR-16-5p | JMJD1C |
| miR-16-5p | FOSL2 |
| miR-16-5p | GMPR |
| miR-16-5p | TFDP1 |
| miR-16-5p | TNFAIP6 |
| miR-16-5p | SIPA1L1 |
| miR-16-5p | WWOX |
| miR-16-5p | HYAL1 |
| miR-16-5p | PSTPIP1 |
| miR-16-5p | CXCL8 |
| miR-16-5p | ADGRL2 |
| miR-16-5p | ACACA |
| miR-16-5p | AOPEP |
| miR-16-5p | PRORP |
| miR-16-5p | IRF1-AS1 |
| miR-223-3p | BCL2 |
| miR-223-3p | VEGFA |
| miR-223-3p | MMP9 |
| miR-223-3p | CAT |
| miR-223-3p | ENTPD1 |
| miR-223-3p | MECP2 |
| miR-223-3p | MMP2 |
| miR-155-5p | BCL2 |
| miR-155-5p | CDK6 |
| miR-155-5p | HSPD1 |
| miR-155-5p | MTX1 |
| miR-155-5p | VEGFA |
| miR-155-5p | FAS |
| miR-155-5p | IL6R |
| miR-155-5p | RERE |
| miR-155-5p | FBN1 |
| miR-155-5p | TLR4 |
| miR-155-5p | CD83 |
| miR-155-5p | CLU |
| miR-155-5p | ABCB1 |
| miR-155-5p | VIM |
| miR-155-5p | CD274 |
| miR-155-5p | CXCR4 |
| miR-155-5p | TP53 |
| miR-155-5p | ELK3 |
| miR-155-5p | NR4A2 |
| miR-155-5p | KLF4 |
| miR-155-5p | MMP1 |
| miR-155-5p | ANXA11 |
| miR-155-5p | RHOBTB1 |
| miR-155-5p | ADA |
| miR-155-5p | CALD1 |
| miR-155-5p | DAPK1 |
| miR-155-5p | DDIT3 |
| miR-155-5p | SLC26A2 |
| miR-155-5p | DUSP1 |
| miR-155-5p | EGR1 |
| miR-155-5p | CTTN |
| miR-155-5p | GAS6 |
| miR-155-5p | GEM |
| miR-155-5p | CXCL1 |
| miR-155-5p | HMOX1 |
| miR-155-5p | IL6ST |
| miR-155-5p | IL7R |
| miR-155-5p | IL18 |
| miR-155-5p | ISG20 |
| miR-155-5p | LIG4 |
| miR-155-5p | LPP |
| miR-155-5p | TNFRSF11B |
| miR-155-5p | PLOD2 |
| miR-155-5p | MAPK1 |
| miR-155-5p | PRNP |
| miR-155-5p | PROS1 |
| miR-155-5p | PTX3 |
| miR-155-5p | SLC2A3 |
| miR-155-5p | TIMP2 |
| miR-155-5p | TNFAIP3 |
| miR-155-5p | UBE2E1 |
| miR-155-5p | USO1 |
| miR-155-5p | ABCC3 |
| miR-155-5p | IL32 |
| miR-155-5p | B4GALT5 |
| miR-155-5p | BMS1 |
| miR-155-5p | SH2B3 |
| miR-155-5p | PLK2 |
| miR-155-5p | ATXN2L |
| miR-155-5p | TNIK |
| miR-155-5p | CRB1 |
| miR-155-5p | MAFF |
| miR-155-5p | ASAP1 |
| miR-155-5p | ETNK1 |
| miR-155-5p | ANKRD9 |
| miR-155-5p | LURAP1L |
| miR-155-5p | BRD2 |
| miR-155-5p | IGF1 |
| miR-155-5p | ADRB2 |
| miR-155-5p | IL4R |
| miR-155-5p | LTA |
| miR-155-5p | IL2RB |
| miR-155-5p | ATN1 |
| miR-155-5p | TNFAIP8 |
| miR-155-5p | POSTN |
| miR-155-5p | TCF7L2 |
| miR-155-5p | EGF |
| miR-155-5p | ATG16L1 |
| miR-155-5p | LRBA |
| miR-155-5p | BMP4 |
| miR-155-5p | HLA-C |
| miR-155-5p | HSPA6 |
| miR-155-5p | GNG11 |
| miR-155-5p | FOSL2 |
| miR-155-5p | HLA-B |
| miR-155-5p | GMPR |
| miR-155-5p | TFDP1 |
| miR-155-5p | TNFAIP6 |
| miR-155-5p | KSR1 |
| miR-155-5p | WWOX |
| miR-155-5p | ADGRL2 |
| miR-155-5p | ACACA |
| miR-155-5p | AOPEP |
| miR-125a-5p | THBS1 |
| miR-125a-5p | JUN |
| miR-125a-5p | IRAK1 |
| miR-125a-5p | NCAM1 |
| miR-125a-5p | CTNNB1 |
| miR-125a-5p | IL6ST |
| miR-125a-5p | PRKAR1A |
| miR-125a-5p | UBE3C |
| miR-125a-5p | TNFSF15 |
| miR-125a-5p | ATXN2L |
| miR-125a-5p | ZFC3H1 |
| miR-125a-5p | TNFRSF1B |
| miR-125a-5p | PTPN2 |
| miR-125a-5p | DUSP4 |
| miR-125a-5p | SIPA1L1 |
| miR-146a-5p | CDK6 |
| miR-146a-5p | THBS1 |
| miR-146a-5p | VEGFA |
| miR-146a-5p | RERE |
| miR-146a-5p | CD83 |
| miR-146a-5p | CARD8 |
| miR-146a-5p | JUN |
| miR-146a-5p | CLU |
| miR-146a-5p | CD274 |
| miR-146a-5p | MMP9 |
| miR-146a-5p | KLF4 |
| miR-146a-5p | FOXP1 |
| miR-146a-5p | ANXA11 |
| miR-146a-5p | CALD1 |
| miR-146a-5p | CD28 |
| miR-146a-5p | CCR4 |
| miR-146a-5p | CXADR |
| miR-146a-5p | SLC26A2 |
| miR-146a-5p | FLNB |
| miR-146a-5p | GAS6 |
| miR-146a-5p | CXCL1 |
| miR-146a-5p | HLA-G |
| miR-146a-5p | IL18 |
| miR-146a-5p | ITGB5 |
| miR-146a-5p | LPP |
| miR-146a-5p | MECP2 |
| miR-146a-5p | MME |
| miR-146a-5p | NOS3 |
| miR-146a-5p | CCL20 |
| miR-146a-5p | SLC2A3 |
| miR-146a-5p | SNAI1 |
| miR-146a-5p | STIM1 |
| miR-146a-5p | STXBP2 |
| miR-146a-5p | TIMP1 |
| miR-146a-5p | TIMP2 |
| miR-146a-5p | TNFAIP3 |
| miR-146a-5p | USO1 |
| miR-146a-5p | ABCC3 |
| miR-146a-5p | IL32 |
| miR-146a-5p | SH2B3 |
| miR-146a-5p | TRIM13 |
| miR-146a-5p | TNIK |
| miR-146a-5p | FNBP1 |
| miR-146a-5p | TTLL5 |
| miR-146a-5p | ASAP1 |
| miR-146a-5p | MEX3C |
| miR-146a-5p | ETNK1 |
| miR-146a-5p | CXCL16 |
| miR-146a-5p | ERAP2 |
| miR-146a-5p | NAA25 |
| miR-146a-5p | SIK1 |
| miR-146a-5p | CXCL10 |
| miR-146a-5p | PSMB9 |
| miR-146a-5p | HFE |
| miR-146a-5p | TWIST1 |
| miR-146a-5p | TNFRSF1B |
| miR-146a-5p | TNFAIP8 |
| miR-146a-5p | FOSB |
| miR-146a-5p | SLC19A1 |
| miR-146a-5p | PTPN2 |
| miR-146a-5p | HLA-DPA1 |
| miR-146a-5p | DUSP4 |
| miR-146a-5p | HLA-C |
| miR-146a-5p | NGF |
| miR-146a-5p | JMJD1C |
| miR-146a-5p | PDCD1 |
| miR-146a-5p | HLA-B |
| miR-146a-5p | GMPR |
| miR-146a-5p | TFDP1 |
| miR-146a-5p | KSR1 |
| miR-146a-5p | SIPA1L1 |
| miR-146a-5p | AOPEP |
| miR-146a-5p | PRORP |
| miR-146a-5p | IRF1-AS1 |
| miR-223-5p | FOS |
| miR-223-5p | SLC2A3 |
| miR-223-5p | ERAP2 |
| miR-223-5p | ATIC |

**Supplementary Table 5.**
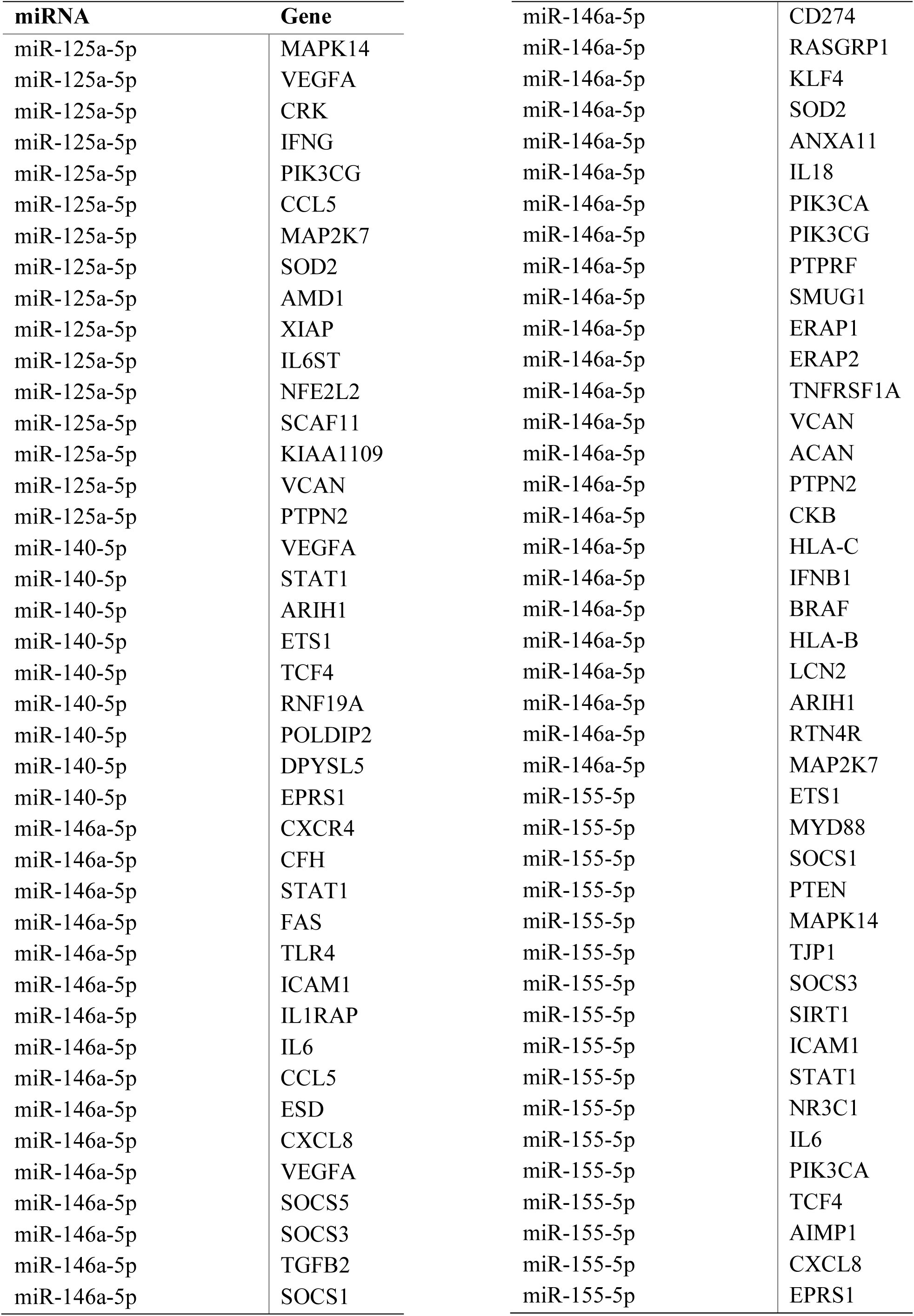
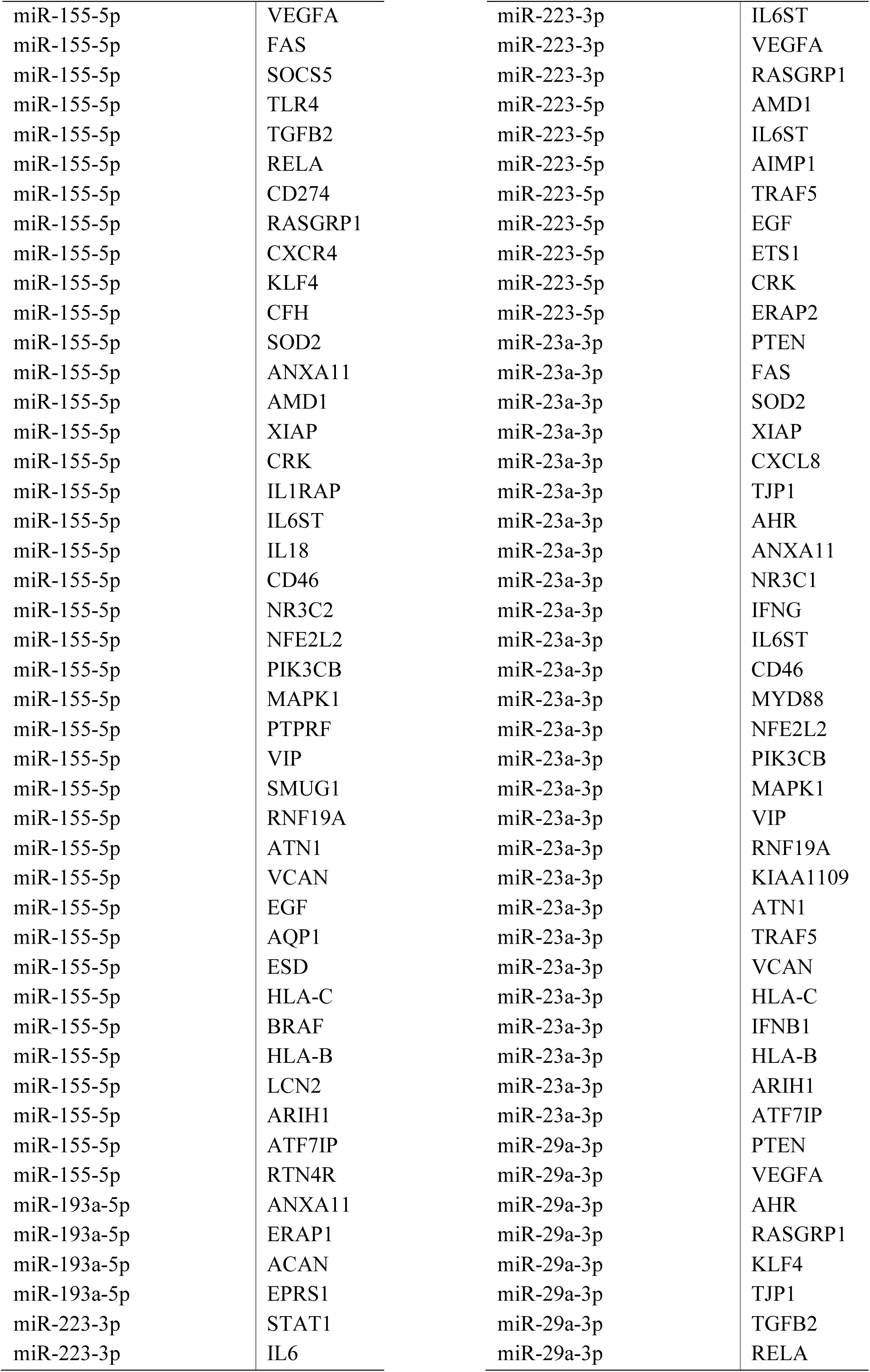
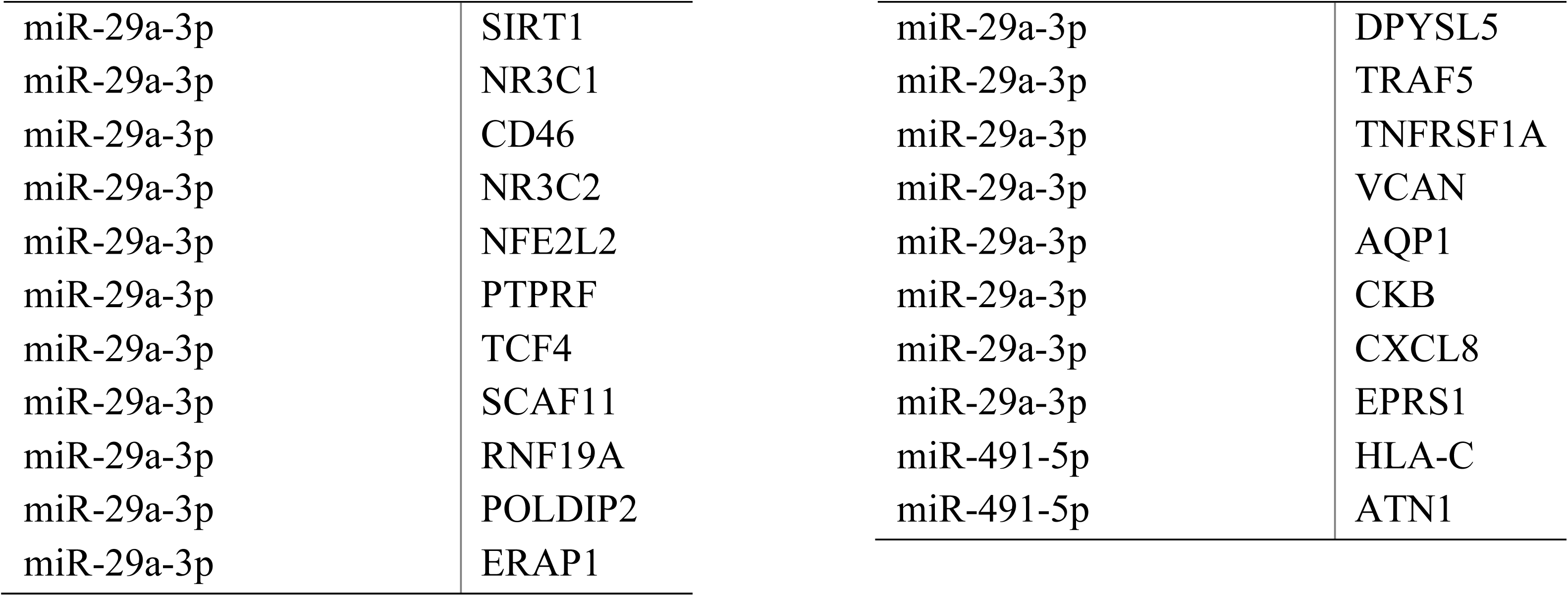
List of genes (74) regulated by more than one of the identified miRNAs in patients uveitis (196 miRNA-gene interaction pairs).

| miRNA | Gene |  |  |
| --- | --- | --- | --- |
| miR-125a-5p | MAPK14 | miR-146a-5p | CD274 |
| miR-125a-5p | VEGFA | miR-146a-5p | RASGRP1 |
| miR-125a-5p | CRK | miR-146a-5p | KLF4 |
| miR-125a-5p | IFNG | miR-146a-5p | SOD2 |
| miR-125a-5p | PIK3CG | miR-146a-5p | ANXA11 |
| miR-125a-5p | CCL5 | miR-146a-5p | IL18 |
| miR-125a-5p | MAP2K7 | miR-146a-5p | PIK3CA |
| miR-125a-5p | SOD2 | miR-146a-5p | PIK3CG |
| miR-125a-5p | AMD1 | miR-146a-5p | PTPRF |
| miR-125a-5p | XIAP | miR-146a-5p | SMUG1 |
| miR-125a-5p | IL6ST | miR-146a-5p | ERAP1 |
| miR-125a-5p | NFE2L2 | miR-146a-5p | ERAP2 |
| miR-125a-5p | SCAF11 | miR-146a-5p | TNFRSF1A |
| miR-125a-5p | KIAA1109 | miR-146a-5p | VCAN |
| miR-125a-5p | VCAN | miR-146a-5p | ACAN |
| miR-125a-5p | PTPN2 | miR-146a-5p | PTPN2 |
| miR-140-5p | VEGFA | miR-146a-5p | CKB |
| miR-140-5p | STAT1 | miR-146a-5p | HLA-C |
| miR-140-5p | ARIH1 | miR-146a-5p | IFNB1 |
| miR-140-5p | ETS1 | miR-146a-5p | BRAF |
| miR-140-5p | TCF4 | miR-146a-5p | HLA-B |
| miR-140-5p | RNF19A | miR-146a-5p | LCN2 |
| miR-140-5p | POLDIP2 | miR-146a-5p | ARIH1 |
| miR-140-5p | DPYSL5 | miR-146a-5p | RTN4R |
| miR-140-5p | EPRS1 | miR-146a-5p | MAP2K7 |
| miR-146a-5p | CXCR4 | miR-155-5p | ETS1 |
| miR-146a-5p | CFH | miR-155-5p | MYD88 |
| miR-146a-5p | STAT1 | miR-155-5p | SOCS1 |
| miR-146a-5p | FAS | miR-155-5p | PTEN |
| miR-146a-5p | TLR4 | miR-155-5p | MAPK14 |
| miR-146a-5p | ICAM1 | miR-155-5p | TJP1 |
| miR-146a-5p | IL1RAP | miR-155-5p | SOCS3 |
| miR-146a-5p | IL6 | miR-155-5p | SIRT1 |
| miR-146a-5p | CCL5 | miR-155-5p | ICAM1 |
| miR-146a-5p | ESD | miR-155-5p | STAT1 |
| miR-146a-5p | CXCL8 | miR-155-5p | NR3C1 |
| miR-146a-5p | VEGFA | miR-155-5p | IL6 |
| miR-146a-5p | SOCS5 | miR-155-5p | PIK3CA |
| miR-146a-5p | SOCS3 | miR-155-5p | TCF4 |
| miR-146a-5p | TGFB2 | miR-155-5p | AIMP1 |
| miR-146a-5p | SOCS1 | miR-155-5p | CXCL8 |
|  |  | miR-155-5p | EPRS1 |

|  |  |
| --- | --- |
| miR-155-5p | VEGFA |
| miR-155-5p | FAS |
| miR-155-5p | SOCS5 |
| miR-155-5p | TLR4 |
| miR-155-5p | TGFB2 |
| miR-155-5p | RELA |
| miR-155-5p | CD274 |
| miR-155-5p | RASGRP1 |
| miR-155-5p | CXCR4 |
| miR-155-5p | KLF4 |
| miR-155-5p | CFH |
| miR-155-5p | SOD2 |
| miR-155-5p | ANXA11 |
| miR-155-5p | AMD1 |
| miR-155-5p | XIAP |
| miR-155-5p | CRK |
| miR-155-5p | IL1RAP |
| miR-155-5p | IL6ST |
| miR-155-5p | IL18 |
| miR-155-5p | CD46 |
| miR-155-5p | NR3C2 |
| miR-155-5p | NFE2L2 |
| miR-155-5p | PIK3CB |
| miR-155-5p | MAPK1 |
| miR-155-5p | PTPRF |
| miR-155-5p | VIP |
| miR-155-5p | SMUG1 |
| miR-155-5p | RNF19A |
| miR-155-5p | ATN1 |
| miR-155-5p | VCAN |
| miR-155-5p | EGF |
| miR-155-5p | AQP1 |
| miR-155-5p | ESD |
| miR-155-5p | HLA-C |
| miR-155-5p | BRAF |
| miR-155-5p | HLA-B |
| miR-155-5p | LCN2 |
| miR-155-5p | ARIH1 |
| miR-155-5p | ATF7IP |
| miR-155-5p | RTN4R |
| miR-193a-5p | ANXA11 |
| miR-193a-5p | ERAP1 |
| miR-193a-5p | ACAN |
| miR-193a-5p | EPRS1 |
| miR-223-3p | STAT1 |
| miR-223-3p | IL6 |
| miR-223-3p | IL6ST |
| miR-223-3p | VEGFA |
| miR-223-3p | RASGRP1 |
| miR-223-5p | AMD1 |
| miR-223-5p | IL6ST |
| miR-223-5p | AIMP1 |
| miR-223-5p | TRAF5 |
| miR-223-5p | EGF |
| miR-223-5p | ETS1 |
| miR-223-5p | CRK |
| miR-223-5p | ERAP2 |
| miR-23a-3p | PTEN |
| miR-23a-3p | FAS |
| miR-23a-3p | SOD2 |
| miR-23a-3p | XIAP |
| miR-23a-3p | CXCL8 |
| miR-23a-3p | TJP1 |
| miR-23a-3p | AHR |
| miR-23a-3p | ANXA11 |
| miR-23a-3p | NR3C1 |
| miR-23a-3p | IFNG |
| miR-23a-3p | IL6ST |
| miR-23a-3p | CD46 |
| miR-23a-3p | MYD88 |
| miR-23a-3p | NFE2L2 |
| miR-23a-3p | PIK3CB |
| miR-23a-3p | MAPK1 |
| miR-23a-3p | VIP |
| miR-23a-3p | RNF19A |
| miR-23a-3p | KIAA1109 |
| miR-23a-3p | ATN1 |
| miR-23a-3p | TRAF5 |
| miR-23a-3p | VCAN |
| miR-23a-3p | HLA-C |
| miR-23a-3p | IFNB1 |
| miR-23a-3p | HLA-B |
| miR-23a-3p | ARIH1 |
| miR-23a-3p | ATF7IP |
| miR-29a-3p | PTEN |
| miR-29a-3p | VEGFA |
| miR-29a-3p | AHR |
| miR-29a-3p | RASGRP1 |
| miR-29a-3p | KLF4 |
| miR-29a-3p | TJP1 |
| miR-29a-3p | TGFB2 |
| miR-29a-3p | RELA |
| miR-29a-3p | SIRT1 |
| miR-29a-3p | NR3C1 |
| miR-29a-3p | CD46 |
| miR-29a-3p | NR3C2 |
| miR-29a-3p | NFE2L2 |
| miR-29a-3p | PTPRF |
| miR-29a-3p | TCF4 |
| miR-29a-3p | SCAF11 |
| miR-29a-3p | RNF19A |
| miR-29a-3p | POLDIP2 |
| miR-29a-3p | ERAP1 |
| miR-29a-3p | DPYSL5 |
| miR-29a-3p | TRAF5 |
| miR-29a-3p | TNFRSF1A |
| miR-29a-3p | VCAN |
| miR-29a-3p | AQP1 |
| miR-29a-3p | CKB |
| miR-29a-3p | CXCL8 |
| miR-29a-3p | EPRS1 |
| miR-491-5p | HLA-C |
| miR-491-5p | ATN1 |

**Supplementary Table 6.**
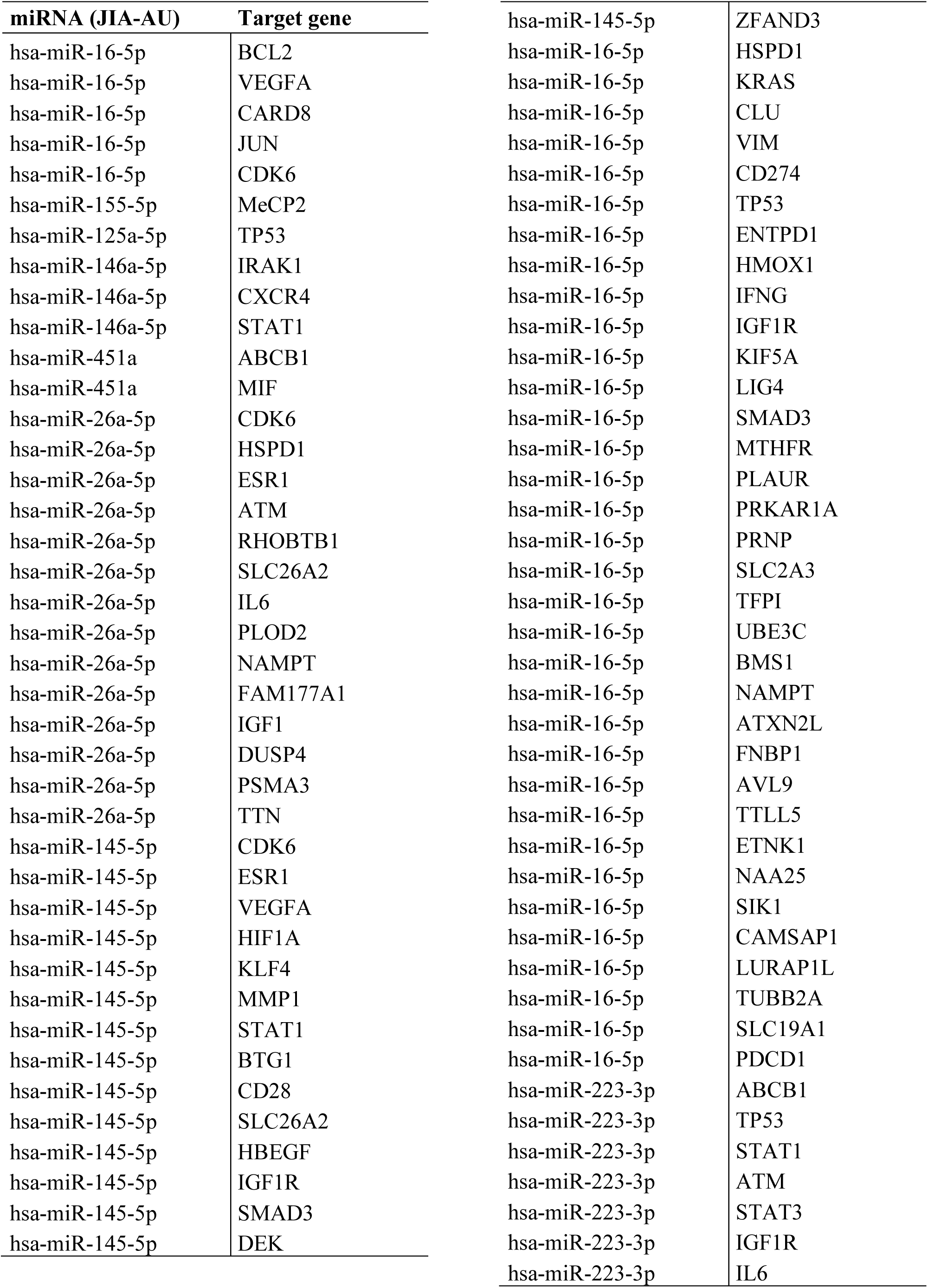
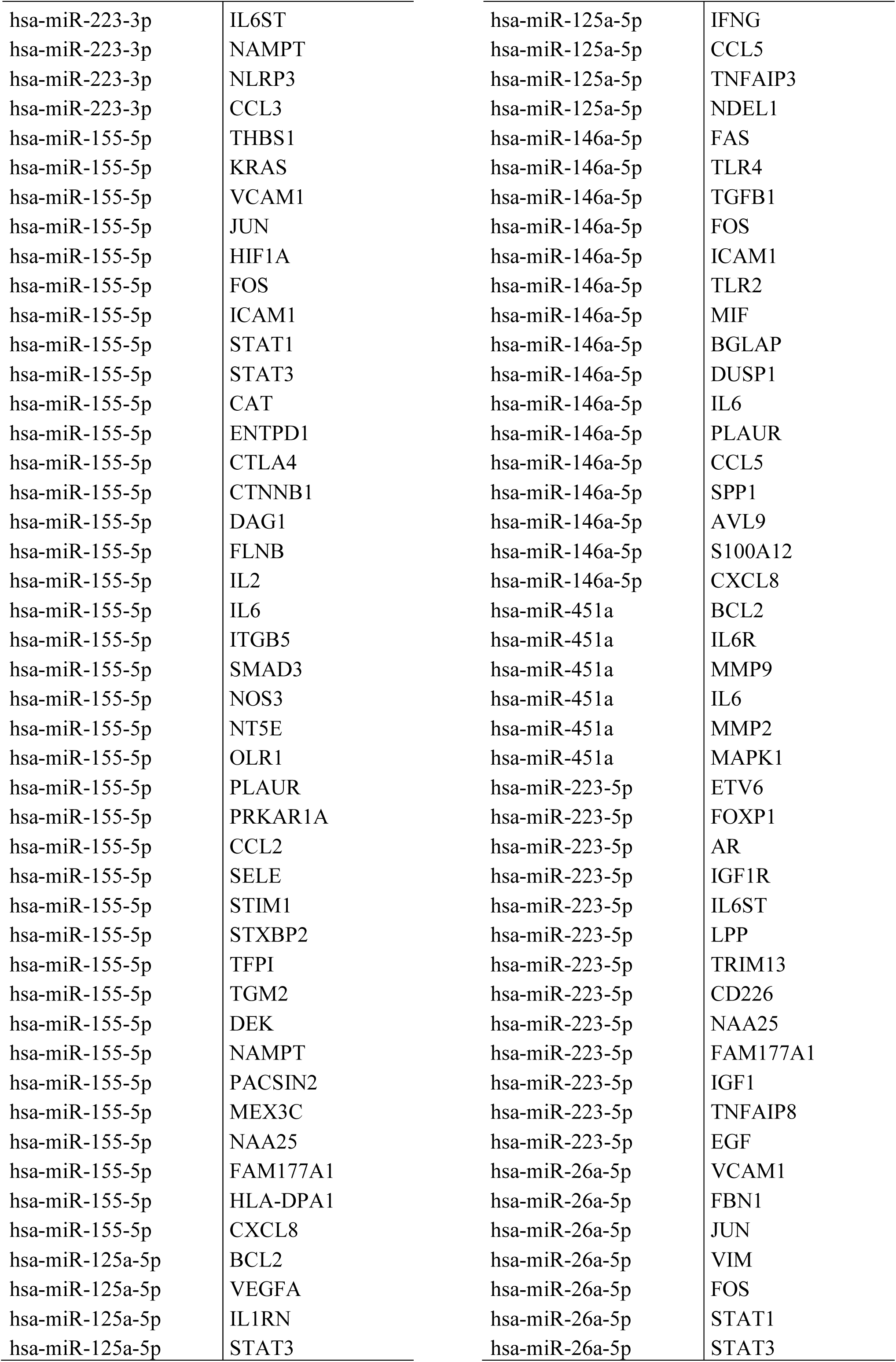
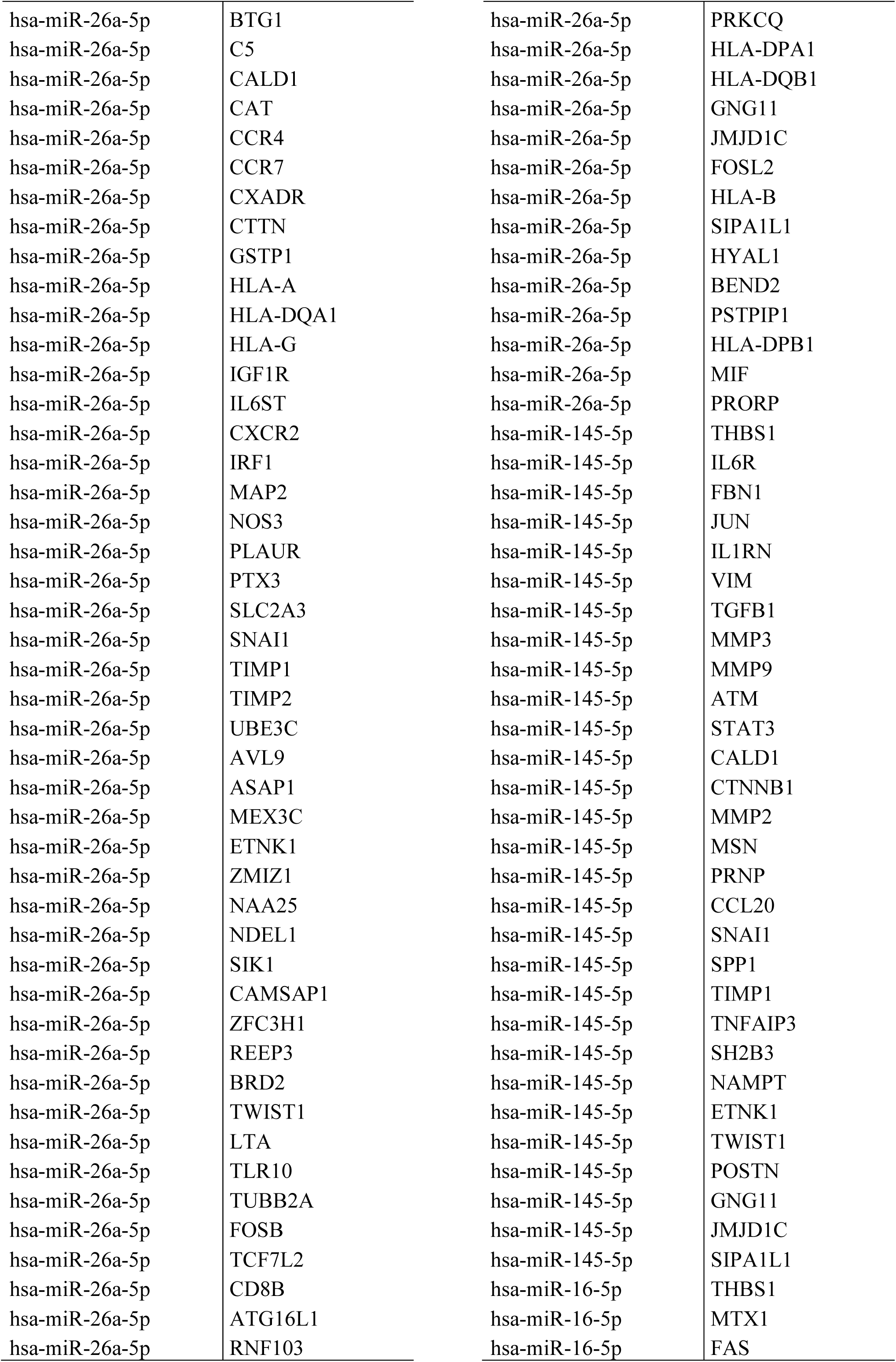
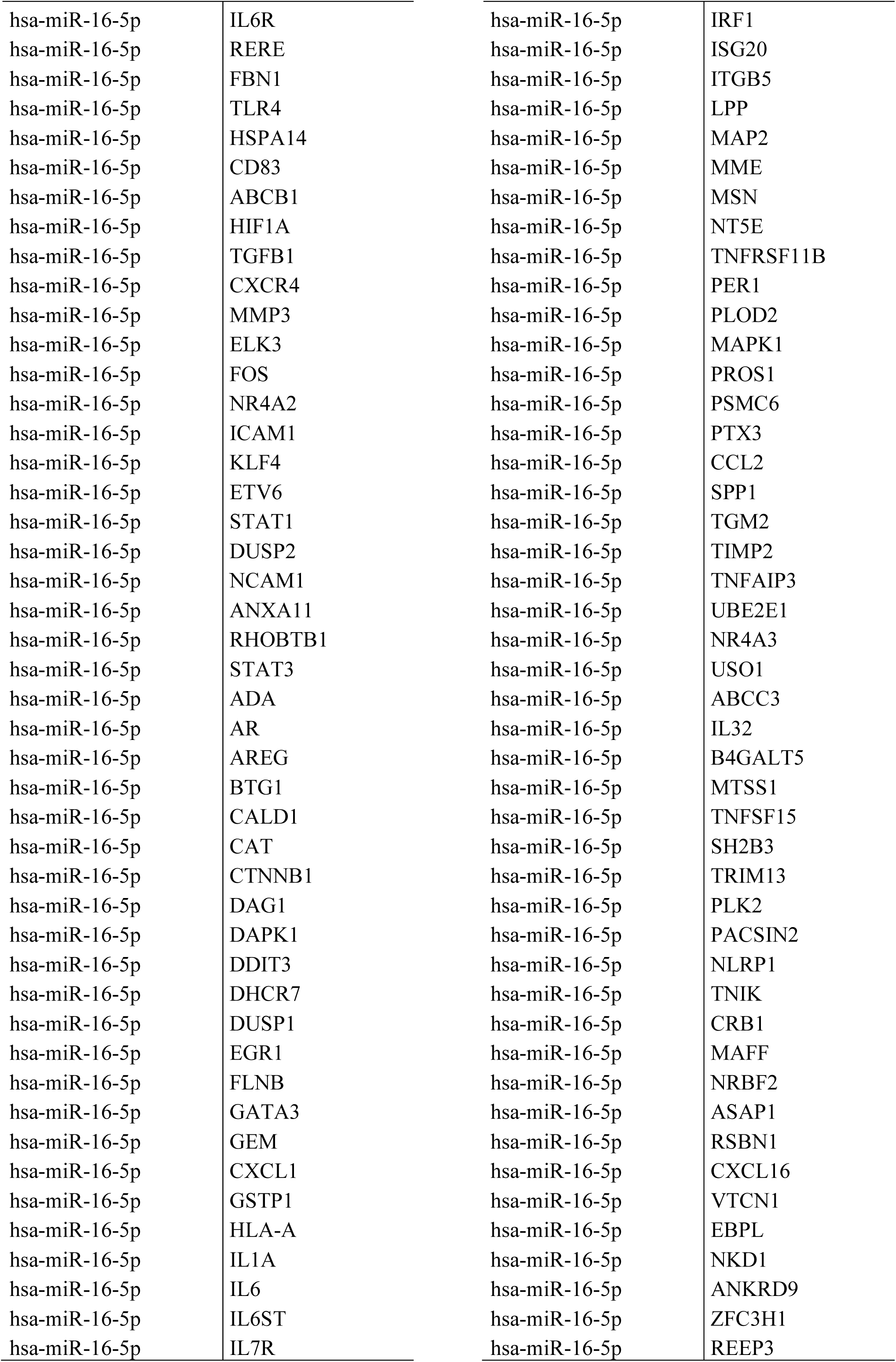
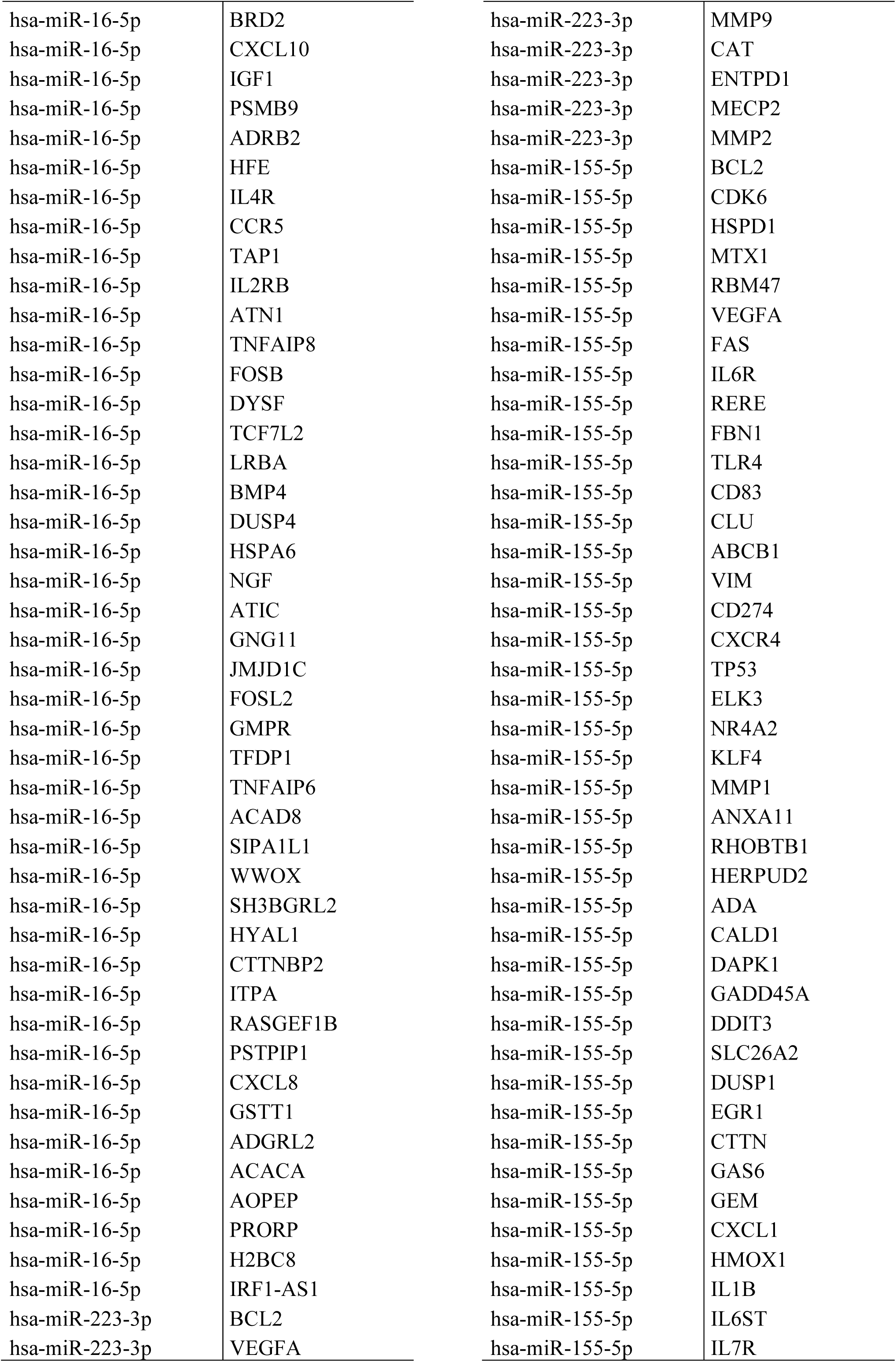
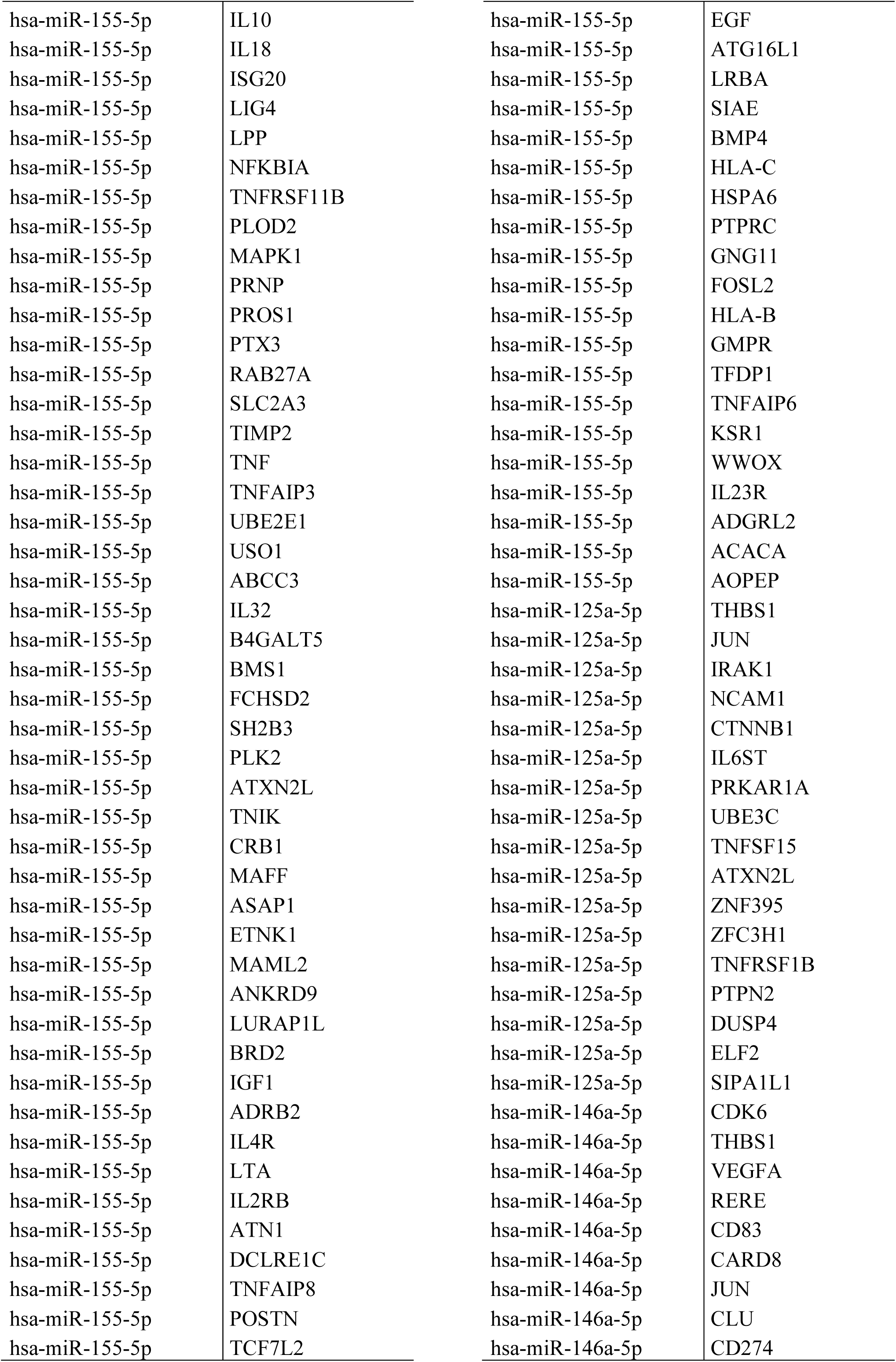
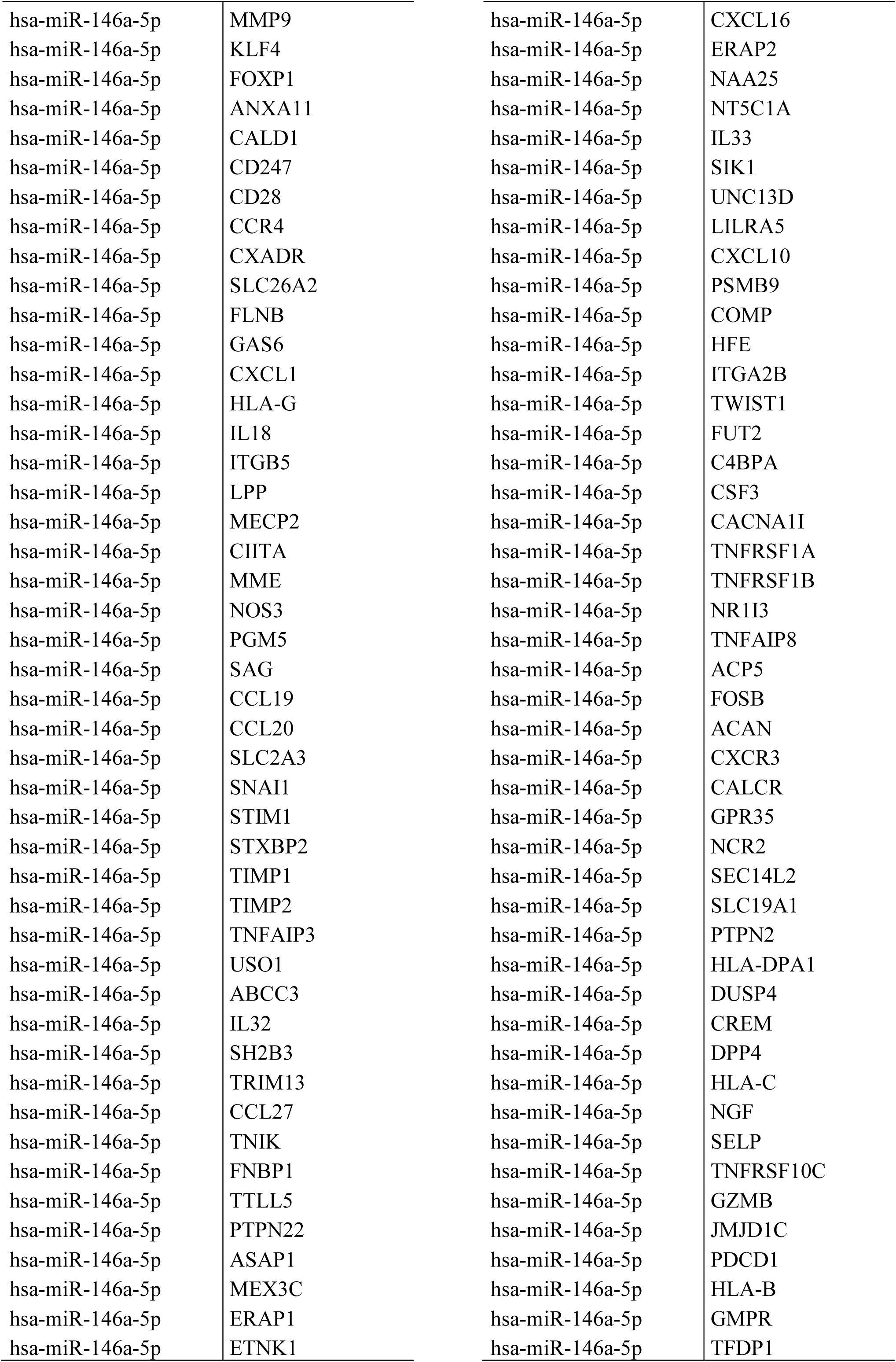
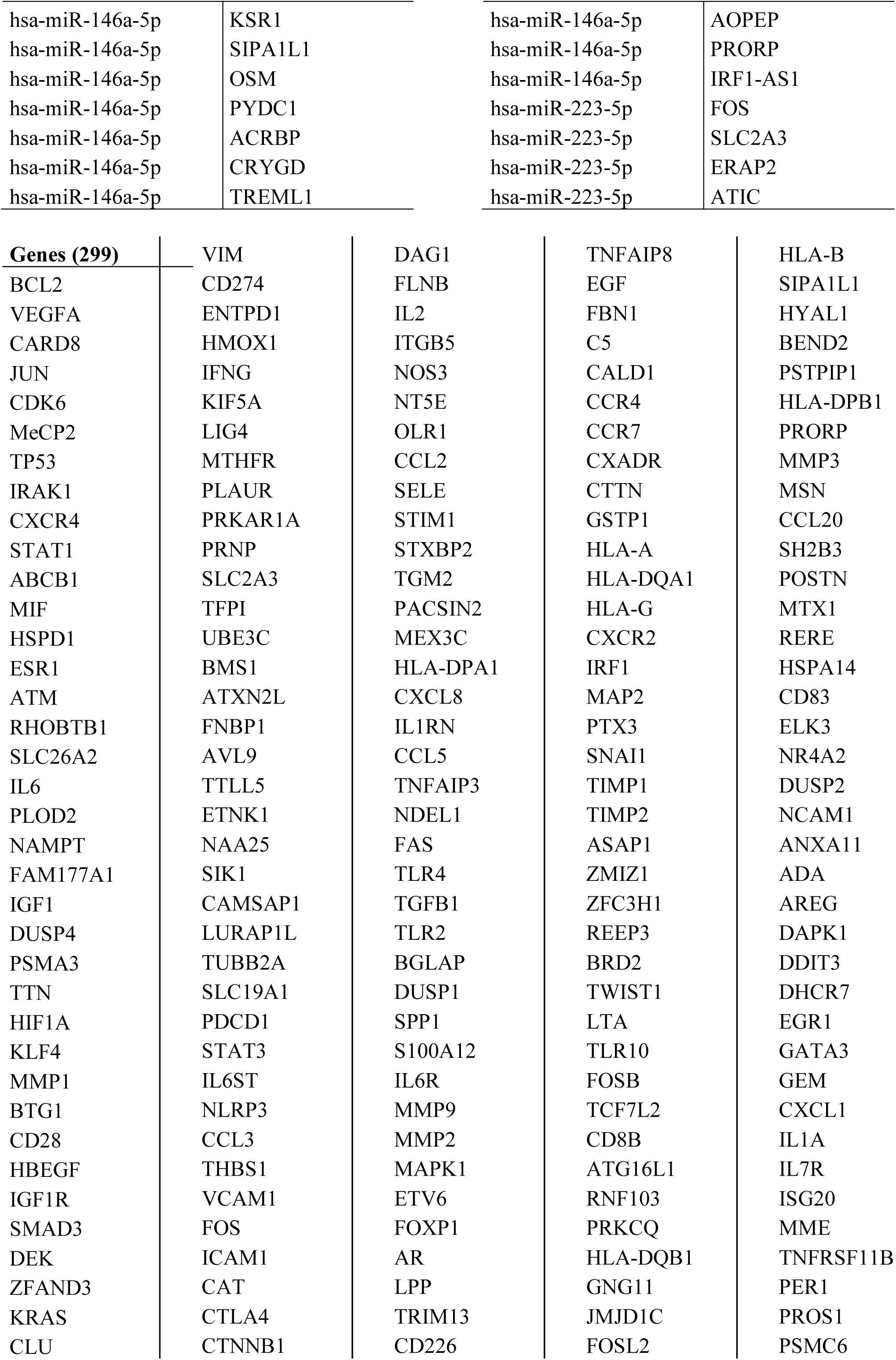
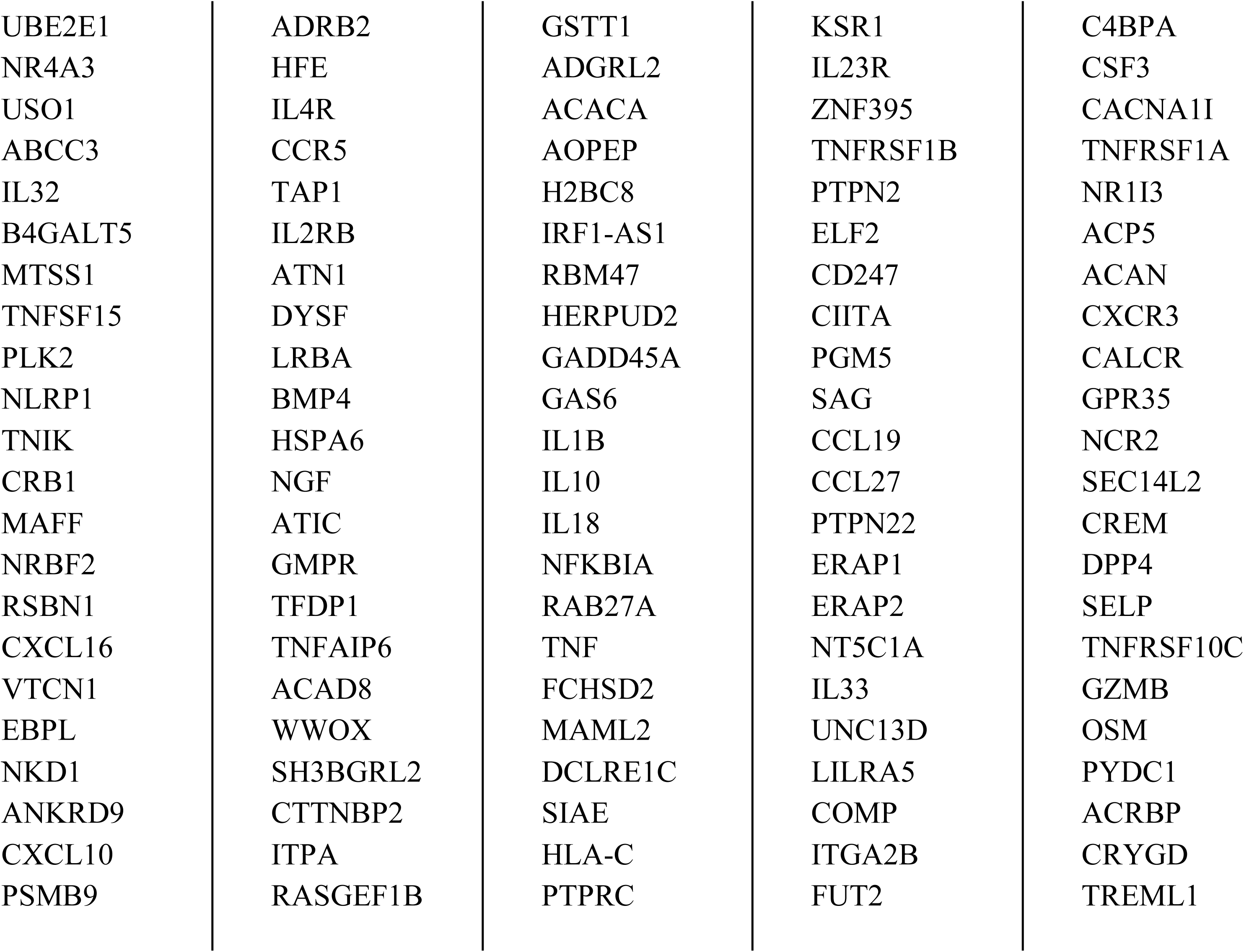
Experimentally validated miRNA:gene pairs found *in silico* among 9 the most frequently detected miRNAs in patients with juvenile arthritis-associated uveitis and 450 genes associated with juvenile arthritis (648 miRNA-gene interaction pairs). From this group, 299 unique genes were selected for pathway enrichment analysis.

**Supplementary Table 7.** Experimentally validated miRNA:gene pairs found *in silico* among 10 the most frequently detected miRNAs in patients with uveitis and 247 genes associated with uveitis (264 miRNA-gene interaction pairs). From this group, 142 unique genes were selected for pathway enrichment analysis.

| miRNA (IU) | Target gene |  |  |
| --- | --- | --- | --- |
| hsa-miR-155-5p | ETS1 | hsa-miR-125a-5p | IL1RN |
| hsa-miR-155-5p | MyD88 | hsa-miR-125a-5p | CRK |
| hsa-miR-155-5p | SOCS1 | hsa-miR-125a-5p | IFNG |
| hsa-miR-140-5p | VEGFA | hsa-miR-125a-5p | PIK3CG |
| hsa-miR-29a-3p | PTEN | hsa-miR-125a-5p | CCL5 |
| hsa-miR-146a-5p | CXCR4 | hsa-miR-125a-5p | MAP2K7 |
| hsa-miR-146a-5p | CFH | hsa-miR-140-5p | STAT1 |
| hsa-miR-146a-5p | STAT1 | hsa-miR-140-5p | ARIH1 |
| hsa-miR-23a-3p | PTEN | hsa-miR-29a-3p | VEGFA |
| hsa-miR-23a-3p | FAS | hsa-miR-29a-3p | AHR |
| hsa-miR-23a-3p | SOD2 | hsa-miR-29a-3p | RASGRP1 |
| hsa-miR-23a-3p | XIAP | hsa-miR-29a-3p | KLF4 |
| hsa-miR-23a-3p | CCL8 | hsa-miR-29a-3p | S100B |
| hsa-miR-23a-3p | CXCL8 | hsa-miR-146a-5p | FAS |
| hsa-miR-223-3p | STAT1 | hsa-miR-146a-5p | TLR4 |
| hsa-miR-223-3p | IL6 | hsa-miR-146a-5p | ICAM1 |
| hsa-miR-223-3p | IL6ST | hsa-miR-146a-5p | TLR2 |
| hsa-miR-223-3p | NLRP3 | hsa-miR-146a-5p | IL1RAP |
| hsa-miR-223-3p | TRPV2 | hsa-miR-146a-5p | IL6 |
| hsa-miR-155-5p | PTEN | hsa-miR-146a-5p | CCL5 |
| hsa-miR-155-5p | MAPK14 | hsa-miR-146a-5p | DECR1 |
| hsa-miR-155-5p | TJP1 | hsa-miR-146a-5p | ESD |
| hsa-miR-155-5p | SOCS3 | hsa-miR-146a-5p | GRAP2 |
| hsa-miR-155-5p | SIRT1 | hsa-miR-146a-5p | S100A12 |
| hsa-miR-155-5p | ICAM1 | hsa-miR-146a-5p | CXCL8 |
| hsa-miR-155-5p | STAT1 | hsa-miR-491-5p | HLA-C |
| hsa-miR-155-5p | CTLA4 | hsa-miR-223-5p | AMD1 |
| hsa-miR-155-5p | NR3C1 | hsa-miR-223-5p | IL6ST |
| hsa-miR-155-5p | IL2 | hsa-miR-223-5p | AIMP1 |
| hsa-miR-155-5p | IL6 | hsa-miR-223-5p | TRAF5 |
| hsa-miR-155-5p | PIK3CA | hsa-miR-223-5p | EGF |
| hsa-miR-155-5p | CCL2 | hsa-miR-23a-3p | TJP1 |
| hsa-miR-155-5p | TCF4 | hsa-miR-23a-3p | HMGB1 |
| hsa-miR-155-5p | AIMP1 | hsa-miR-23a-3p | AHR |
| hsa-miR-155-5p | SLC9A3R2 | hsa-miR-23a-3p | TGFBR3 |
| hsa-miR-155-5p | CXCL8 | hsa-miR-23a-3p | ANXA11 |
| hsa-miR-155-5p | EPRS1 | hsa-miR-23a-3p | NR3C1 |
| hsa-miR-125a-5p | MAPK14 | hsa-miR-23a-3p | HLA-A |
| hsa-miR-125a-5p | ZEB2 | hsa-miR-23a-3p | IFNG |
| hsa-miR-125a-5p | VEGFA | hsa-miR-23a-3p | IL6ST |
|  |  | hsa-miR-23a-3p | CD46 |
|  |  | hsa-miR-23a-3p | MYD88 |
| hsa-miR-23a-3p | NFE2L2 | hsa-miR-155-5p | IL15 |
| hsa-miR-23a-3p | PIK3CB | hsa-miR-155-5p | IL18 |
| hsa-miR-23a-3p | MAPK1 | hsa-miR-155-5p | CD46 |
| hsa-miR-23a-3p | VIP | hsa-miR-155-5p | NR3C2 |
| hsa-miR-23a-3p | RNF19A | hsa-miR-155-5p | NFE2L2 |
| hsa-miR-23a-3p | KIAA1109 | hsa-miR-155-5p | PIK3CB |
| hsa-miR-23a-3p | ATN1 | hsa-miR-155-5p | PLXNA2 |
| hsa-miR-23a-3p | TRAF5 | hsa-miR-155-5p | MAPK1 |
| hsa-miR-23a-3p | VCAN | hsa-miR-155-5p | PTPRF |
| hsa-miR-23a-3p | AQP4 | hsa-miR-155-5p | TNF |
| hsa-miR-23a-3p | HLA-C | hsa-miR-155-5p | VIP |
| hsa-miR-23a-3p | IFNB1 | hsa-miR-155-5p | AHSA1 |
| hsa-miR-23a-3p | HLA-B | hsa-miR-155-5p | SMUG1 |
| hsa-miR-23a-3p | ARIH1 | hsa-miR-155-5p | RNF19A |
| hsa-miR-23a-3p | ATF7IP | hsa-miR-155-5p | ALB |
| hsa-miR-23a-3p | DNER | hsa-miR-155-5p | ATN1 |
| hsa-miR-223-3p | VEGFA | hsa-miR-155-5p | VCAN |
| hsa-miR-223-3p | RASGRP1 | hsa-miR-155-5p | ANGPT1 |
| hsa-miR-223-3p | PIK3CD | hsa-miR-155-5p | EGF |
| hsa-miR-155-5p | MTX1 | hsa-miR-155-5p | AQP1 |
| hsa-miR-155-5p | VEGFA | hsa-miR-155-5p | ESD |
| hsa-miR-155-5p | FAS | hsa-miR-155-5p | HLA-C |
| hsa-miR-155-5p | SOCS5 | hsa-miR-155-5p | EPCAM |
| hsa-miR-155-5p | TLR4 | hsa-miR-155-5p | BRAF |
| hsa-miR-155-5p | TGFB2 | hsa-miR-155-5p | HLA-B |
| hsa-miR-155-5p | RELA | hsa-miR-155-5p | LCN2 |
| hsa-miR-155-5p | ANXA1 | hsa-miR-155-5p | ARIH1 |
| hsa-miR-155-5p | CD274 | hsa-miR-155-5p | PRKCD |
| hsa-miR-155-5p | RASGRP1 | hsa-miR-155-5p | PLA2G15 |
| hsa-miR-155-5p | CXCR4 | hsa-miR-155-5p | MBTPS2 |
| hsa-miR-155-5p | EGR2 | hsa-miR-155-5p | ATF7IP |
| hsa-miR-155-5p | KLF4 | hsa-miR-155-5p | RTN4R |
| hsa-miR-155-5p | CFH | hsa-miR-155-5p | IL17D |
| hsa-miR-155-5p | SOD2 | hsa-miR-155-5p | IL23R |
| hsa-miR-155-5p | ANXA11 | hsa-miR-125a-5p | P2RX7 |
| hsa-miR-155-5p | ADM | hsa-miR-125a-5p | SOD2 |
| hsa-miR-155-5p | AMD1 | hsa-miR-125a-5p | AMD1 |
| hsa-miR-155-5p | XIAP | hsa-miR-125a-5p | XIAP |
| hsa-miR-155-5p | CRK | hsa-miR-125a-5p | IL6ST |
| hsa-miR-155-5p | CYP24A1 | hsa-miR-125a-5p | NFE2L2 |
| hsa-miR-155-5p | GEM | hsa-miR-125a-5p | SCAF11 |
| hsa-miR-155-5p | HMOX1 | hsa-miR-125a-5p | KIAA1109 |
| hsa-miR-155-5p | IL1B | hsa-miR-125a-5p | VCAN |
| hsa-miR-155-5p | IL1RAP | hsa-miR-125a-5p | PTPN2 |
| hsa-miR-155-5p | IL6ST | hsa-miR-125a-5p | MAP6 |
| hsa-miR-155-5p | IL10 | hsa-miR-140-5p | ETS1 |
| hsa-miR-140-5p | TCF4 | hsa-miR-146a-5p | CYP27B1 |
| hsa-miR-140-5p | RNF19A | hsa-miR-146a-5p | CFI |
| hsa-miR-140-5p | POLDIP2 | hsa-miR-146a-5p | IL18 |
| hsa-miR-140-5p | DPYSL5 | hsa-miR-146a-5p | NOS2 |
| hsa-miR-140-5p | EPRS1 | hsa-miR-146a-5p | PIK3CA |
| hsa-miR-29a-3p | TJP1 | hsa-miR-146a-5p | PIK3CG |
| hsa-miR-29a-3p | TGFB2 | hsa-miR-146a-5p | MAPK3 |
| hsa-miR-29a-3p | RELA | hsa-miR-146a-5p | PTPRF |
| hsa-miR-29a-3p | SIRT1 | hsa-miR-146a-5p | SAG |
| hsa-miR-29a-3p | AKR1B1 | hsa-miR-146a-5p | CCL20 |
| hsa-miR-29a-3p | EXTL3 | hsa-miR-146a-5p | TM7SF2 |
| hsa-miR-29a-3p | NR3C1 | hsa-miR-146a-5p | SMUG1 |
| hsa-miR-29a-3p | CD46 | hsa-miR-146a-5p | ERAP1 |
| hsa-miR-29a-3p | NR3C2 | hsa-miR-146a-5p | CXCL16 |
| hsa-miR-29a-3p | NFE2L2 | hsa-miR-146a-5p | NLRC4 |
| hsa-miR-29a-3p | PTPRF | hsa-miR-146a-5p | ERAP2 |
| hsa-miR-29a-3p | TCF4 | hsa-miR-146a-5p | CXCL10 |
| hsa-miR-29a-3p | AIMP2 | hsa-miR-146a-5p | PSMB9 |
| hsa-miR-29a-3p | SCAF11 | hsa-miR-146a-5p | TNFRSF1A |
| hsa-miR-29a-3p | RNF19A | hsa-miR-146a-5p | VCAN |
| hsa-miR-29a-3p | POLDIP2 | hsa-miR-146a-5p | ACAN |
| hsa-miR-29a-3p | TRIB2 | hsa-miR-146a-5p | GPR35 |
| hsa-miR-29a-3p | ERAP1 | hsa-miR-146a-5p | PTPN2 |
| hsa-miR-29a-3p | DPYSL5 | hsa-miR-146a-5p | CD40 |
| hsa-miR-29a-3p | ADO | hsa-miR-146a-5p | AIF1 |
| hsa-miR-29a-3p | TRAF5 | hsa-miR-146a-5p | CKB |
| hsa-miR-29a-3p | TNFRSF1A | hsa-miR-146a-5p | HLA-C |
| hsa-miR-29a-3p | VCAN | hsa-miR-146a-5p | IFNB1 |
| hsa-miR-29a-3p | AQP1 | hsa-miR-146a-5p | BRAF |
| hsa-miR-29a-3p | CKB | hsa-miR-146a-5p | PDCD1 |
| hsa-miR-29a-3p | CXCL8 | hsa-miR-146a-5p | HLA-B |
| hsa-miR-29a-3p | EPRS1 | hsa-miR-146a-5p | LCN2 |
| hsa-miR-146a-5p | VEGFA | hsa-miR-146a-5p | ARIH1 |
| hsa-miR-146a-5p | ICOS | hsa-miR-146a-5p | RTN4R |
| hsa-miR-146a-5p | SOCS5 | hsa-miR-146a-5p | MAP2K7 |
| hsa-miR-146a-5p | SOCS3 | hsa-miR-146a-5p | MZB1 |
| hsa-miR-146a-5p | TGFB2 | hsa-miR-146a-5p | IFNL2 |
| hsa-miR-146a-5p | SOCS1 | hsa-miR-491-5p | ATN1 |
| hsa-miR-146a-5p | ASS1 | hsa-miR-193a-5p | ANXA11 |
| hsa-miR-146a-5p | CD274 | hsa-miR-193a-5p | ERAP1 |
| hsa-miR-146a-5p | RASGRP1 | hsa-miR-193a-5p | ACAN |
| hsa-miR-146a-5p | KLF4 | hsa-miR-193a-5p | EPRS1 |
| hsa-miR-146a-5p | SOD2 | hsa-miR-223-5p | ETS1 |
| hsa-miR-146a-5p | ANXA11 | hsa-miR-223-5p | CRK |
| hsa-miR-146a-5p | CRYAA | hsa-miR-223-5p | ERAP2 |

| Genes (142) |  |  |  |  |
| --- | --- | --- | --- | --- |
| ETS1 | AIMP1 | HLA-A | IL15 | TRIB2 |
| MyD88 | SLC9A3R2 | CD46 | IL18 | ERAP1 |
| SOCS1 | EPRS1 | NFE2L2 | NR3C2 | ADO |
| VEGFA | ZEB2 | PIK3CB | PLXNA2 | TNFRSF1A |
| PTEN | IL1RN | MAPK1 | PTPRF | CKB |
| CXCR4 | CRK | VIP | TNF | ICOS |
| CFH | IFNG | RNF19A | AHSA1 | ASS1 |
| STAT1 | PIK3CG | KIAA1109 | SMUG1 | CRYAA |
| FAS | CCL5 | ATN1 | ALB | CYP27B1 |
| SOD2 | MAP2K7 | VCAN | ANGPT1 | CFI |
| XIAP | ARIH1 | AQP4 | AQP1 | NOS2 |
| CCL8 | AHR | IFNB1 | EPCAM | MAPK3 |
| CXCL8 | RASGRP1 | HLA-B | BRAF | SAG |
| IL6 | KLF4 | ATF7IP | LCN2 | CCL20 |
| IL6ST | S100B | DNER | PRKCD | TM7SF2 |
| NLRP3 | TLR4 | PIK3CD | PLA2G15 | CXCL16 |
| TRPV2 | TLR2 | MTX1 | MBTPS2 | NLRC4 |
| MAPK14 | IL1RAP | SOCS5 | RTN4R | ERAP2 |
| TJP1 | DECR1 | TGFB2 | IL17D | CXCL10 |
| SOCS3 | ESD | RELA | IL23R | PSMB9 |
| SIRT1 | GRAP2 | ANXA1 | P2RX7 | ACAN |
| ICAM1 | S100A12 | CD274 | SCAF11 | GPR35 |
| CTLA4 | HLA-C | EGR2 | PTPN2 | CD40 |
| NR3C1 | AMD1 | ADM | MAP6 | AIF1 |
| IL2 | TRAF5 | CYP24A1 | POLDIP2 | PDCD1 |
| PIK3CA | EGF | GEM | DPYSL5 | MZB1 |
| CCL2 | HMGB1 | HMOX1 | AKR1B1 | IFNL2 |
| TCF4 | TGFBR3 | IL1B | EXTL3 |  |
|  | ANXA11 | IL10 | AIMP2 |  |

